# Food waste as a resource: grinding, dilution, and storage as a pretreatment strategy to produce fermentation intermediates

**DOI:** 10.1101/2020.04.27.064808

**Authors:** Sarah E. Daly, Joseph G. Usack, Lauren A. Harroff, James G. Booth, Michael P. Keleman, Largus T. Angenent

## Abstract

In several states of the U.S., one measure to mitigate greenhouse-gas emissions has been to ban food wastes from landfills. As a result, U.S.-based companies are now providing decentralized food-waste management systems for supermarkets and restaurants, which include storage as a slurry. It is unclear, however, which storage conditions (factors) would affect the spontaneous microbial activity, resulting in a different fermentation product spectra, and how this would affect further post-treatment. Here, we performed two experiments to mimic: 1) storage and 2) subsequent anaerobic digestion. For the food-waste storage system, we designed a mixed-level fractional factorial analysis with 12 experimental combinations, including separating food waste into: carbohydrate-rich, lipid-rich, and protein-rich food waste. We found that all factors that we tested correlated with the fermentation product spectra, but that especially the factors: i) storage temperature; ii) food-waste composition; and iii) storage time affected the fermentation outcome. We observed that relatively low pH levels of 3-4, which were achieved due to rapid lactic acid accumulation by microbial activity during storage, coincided with greater lactate production at a maximum chemical oxygen demand (COD) selectivity of 90%. This provides an opportunity to optimize lactate production, which is ideal for subsequent methane or chemical production.

**TOC/Abstract graphic:** 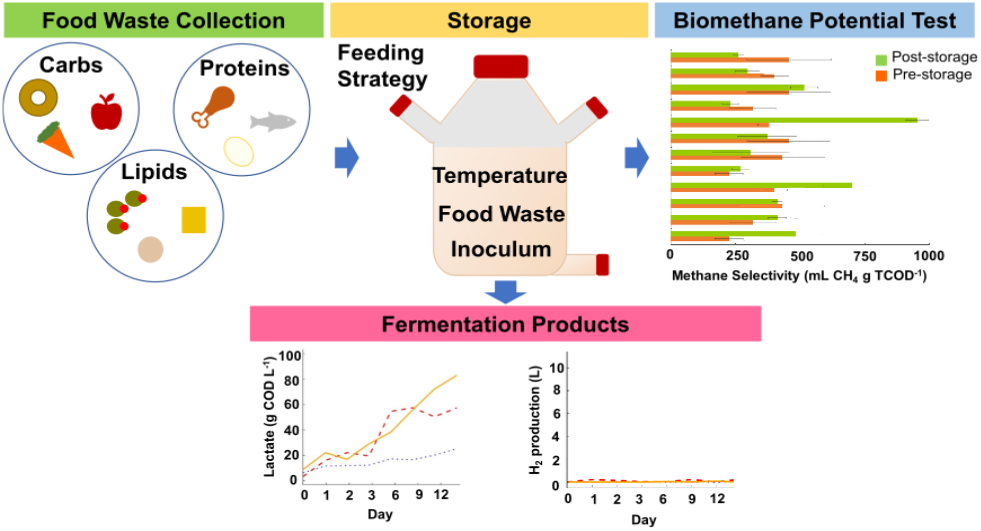

## Introduction

In 2014 alone, U.S. residents produced 38.4×10^6^ tons of food waste with ∼75% going to landfills ^1^. In landfills, these organic wastes are partly converted under anaerobic conditions into methane, which is a potent greenhouse gas (GHG) when released to the atmosphere. The methane release by landfills (without gas capture) contributed 20% of anthropogenic methane emissions in the U.S., which is the third largest methane source in the U.S. ^2^. Since food waste represents ∼40% of organic material going to the landfill ^2^, it is an important anthropogenic source of U.S. methane emissions. On the other hand, the energy content of food waste is estimated to be ∼2% of annual U.S. energy consumption ^3^. Through resource recovery of food waste, considerable amounts of various energy carriers, such as methane *via* anaerobic digestion (AD), can be produced. Then, GHG emissions can be mitigated by reducing methane release to the atmosphere from landfills and offsetting fossil fuel derived carbon-dioxide release from offsetting natural gas consumption. While the energy content of food waste cannot supply the current energy demand, the mitigation in GHG release will be substantial, which is why six U.S. states have already banned landfilling of food waste.

Food waste is an organic source for AD due to its high biodegradability, methane yield, and organic carbon content ^4,5^, especially when used during co-digestion ^6^, which is allowed in many states of the U.S., including New York State. Besides many advantages of co-digestion, it is important to understand that co-digestion of food waste does result in additional nutrient loading, which may have negative environmental conditions ^7^. AD is both reliable and economically feasible at full-scale and is one of the best treatments for agro-industrial, agricultural, and food waste due to its ability to stabilize wastes, while at the same time recover energy ^8–11^.

The simultaneous ban on landfilling food waste and the realization that resource recovery is necessary, has led U.S.-based companies to develop commercial food-waste management systems. Much of the food waste is generated in a decentralized manner at, for example, restaurants and supermarkets, without specialized waste-management personnel. In addition, food waste in temporarily stored bins will decompose and create smells or attract insects, rodents, and even larger animals. Their management system promises to provide a solution to restaurants and supermarkets. Often these systems include the simultaneous grinding and dilution of food waste in the kitchen area, resulting in a slurry. This slurry is then pumped and stored in a closed storage tank, which can be placed inside or outside of the facility. After a one- to two-week storage period, local contractors transport the slurry to the AD facility.

During storage at the restaurant or supermarket site, the food waste undergoes rapid hydrolysis and acidification due to microbial activity as a batch-fermentation system under anaerobic conditions. The storage tank is periodically filled with food-waste slurry during the one- to two-week storage period. Food waste fermentation is spontaneous with indigenous microbes, and thus does not require external inoculum. This was shown by several studies with the goal to specifically convert food waste into hydrogen gas ^12–14^. Their experiments were performed in bottles within a period of several days, and showed that homofermentation to only lactate and/or heterofermentation to acetate, lactate, carbon dioxide, and ethanol ^15^ should be prevented to promote saccharolytic clostridial fermentation ^16,17^ to *n*-butyrate, carbon dioxide, and hydrogen.

It is, however, easier to promote lactate production than hydrogen production, and this was shown with additional experiments and indigenous microbes in food waste ^18,19^. A very recent study with food waste included a control without inoculation, but included an initial pH control to promote heterofermentation ^20^. These results from fermentation of food waste with indigenous microbes (*i.e*., no inoculation) toward lactate production seem to follow the apparent fermentation succession during ensilage ^16,21^, which is also clearly described in a recent co-ensilage study with sugar beet leaves and lignocellulosic material at controlled conditions ^22^. The succession without oxygen is often started by homofermentation and possibly heterofermentation when the pH is not lowered to 4.5 quickly enough ^23^. If the pH remains higher than 4.5 during the initial stages, lactic acid is further converted by saccharolytic clostridial fermentation, which is then a secondary fermentation, into a mixture of carboxylic acids and hydrogen ^16,17^ Only after a very long operating period of months without oxygen is some methane formed when pH levels are still too high ^22^.

However, during food-waste storage the environmental conditions are not controlled, are different depending on season and location, and may vary throughout the storage period. None of the above-mentioned studies with food waste provided an elaborate experiment to include many different factors, which would also be interdependent. As a result, many questions were raised on how storage conditions (factors) could change the fermentation outcome (product spectra) for food waste after diluting and grinding. In addition, questions remained on how further treatment with AD would behave with the different fermentation products. To answer these questions, we performed: 1) a **storage experiment** with large-media bottles in a fractional factorial design; and 2) a **biological-methane-potential experiment** with small-media bottles to determine the change in methane potential of the food waste after an operating period of 15 days.

## Materials and methods

### Storage experiment

#### Experimental design

For the storage experiment, we performed a controlled, large-media-bottle experiment (working volume of 3.75 L; **Fig. S1** and **Supporting Information**), which accounted for the four experimental factors (storage temperature, food-waste composition, feeding strategy, and inoculation) *plus* time in a comprehensive and statistically meaningful way. Because multiple test conditions (two or three levels) were required for each of these four experimental factors, the use of a standard, full-factorial experimental design was not feasible (*i.e*., 3×3×3×2 = 54 experimental combinations), especially when triplicate biological experiments are required (162 large-media bottles). Instead, we used a mixed-level fractional factorial analysis (MFFA) to reduce the total number of experimental combinations by excluding redundant interactions between factors (**Table 1**) ^24–26^. The term “mixed-level” signifies that the factors did not always involve an equal number of test conditions. For example, we tested three storage temperature conditions (*i.e*., 15°C, 25°C, and 35°C), while we only needed two inoculum conditions (*i.e*., inoculum present *vs*. inoculum not present). After defining the factors and the corresponding test conditions, we used AlgDesign in *R* to create the MFFA design using the Federov exchange algorithm (R Studio *v*3.2.2, The R Foundation for Statistical Computing Platform). The program produced 12 unique, experimental combinations, which we then ran in triplicate (36 large-media bottles) (**Table 1**), which resulted in 6 operating periods with 6 large serum bottles for each period.

**Table 1.** Twelve experimental combinations. One test condition from each of the four factors was used to define a unique combination for each storage experiment. The four factors were: storage temperature, food-waste composition, feeding strategy, and the presence of an inoculum. In respect to these factors, we tested three storage temperature conditions: 15°C, 25°C, and 35°C; three food-waste compositions: carbohydrate-rich, lipid-rich, and protein-rich; three feeding strategies: batch, semi-batch at semi-oxic conditions, and semi-batch at anoxic conditions; and two inoculation conditions: with and without inoculum.

| Combination | Storage temperature | Food-waste composition | Feeding strategy | Inoculation |
| --- | --- | --- | --- | --- |
| 1 | 15°C | Carbohydrate-rich | Semi-Batch: Semi-oxic | Yes |
| 2 | 15°C | Carbohydrate-rich | Semi-Batch: Semi-oxic | No |
| 3 | 15°C | Lipid-rich | Batch | No |
| 4 | 15°C | Protein-rich | Semi-Batch: Anoxic | Yes |
| 5 | 25°C | Carbohydrate-rich | Batch | Yes |
| 6 | 25°C | Lipid-rich | Batch | No |
| 7 | 25°C | Lipid-rich | Semi-Batch: Anoxic | Yes |
| 8 | 25°C | Protein-rich | Semi-Batch: Semi-oxic | No |
| 9 | 35°C | Carbohydrate-rich | Semi-Batch: Anoxic | No |
| 10 | 35°C | Lipid-rich | Semi-Batch: Semi-oxic | Yes |
| 11 | 35°C | Protein-rich | Batch | Yes |
| 12 | 35°C | Protein-rich | Semi-Batch: Anoxic | No |

We included 288 sampling points because of the 12 experimental combinations x 3 [triplicates] x 8 sampling points throughout the operating period of 15 days (Days 0, 1, 2, 3, 6, 9, 12, and 15). With the entire dataset, we utilized the multivariate analysis of variance (MANOVA) in the stats package in R Studio *v3.4.4* to calculate which of the different factors *plus* time in the experiment or their two-way and three-way interactions influenced the dependent variables (dot plot visualization). Again with the entire dataset, we utilized principle component analysis (PCA) to visualize the clustering of factors in the data with PCA plots and to determine which dependent variables clustered together in a PCA bi-plot, using the FactoMineR and factoextra packages in R Studio *v*3.4.4. Finally, we utilized the interaction regression model method as described by Kutner et al. ^27^ in paragraph 8.2 with a subset of the dataset (for each of the three food-waste compositions) to develop the interaction plots. We developed the interaction plots for two factors for each dependent variable, using the stats package in R Studio *v*3.4.4. With two factors, each of the interaction plots identifies the interactions between two factors in linear regression models with:

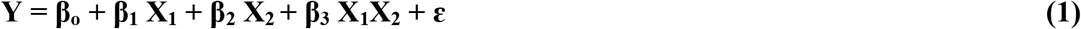

where **Y:** Dependent variable, **β_o_:** Intercept, **X_1_:** Factor 1, **X_2_:** Factor 2, **X_1_X_2_:** Interaction between Factor 1 and 2, and **ε:** Residuals. Finally, the linear regression equations (**Eq. S1-S2**) were developed, using RegBest in FactoMine R (R Studio *v*3.4.4).

#### Food-waste collection and inoculum

We collected pre-consumer food waste from a dining hall at Cornell University, and we manually sorted the food waste by food item. We consulted the USDA Food Composition Database to categorize each food item as either: carbohydrate-rich, lipid-rich, or protein-rich, which was based on the component with the largest mass fraction (*w/w*) ^28^ (**Table 2**). We excluded any item that could not be easily identified. Next, we ground each batch of food waste separately using an InSinkerator® food disposal system (Model MSLV-7, Emerson Electric Co, Racine, WI). The maximum particle size of the ground food waste was ∼5 mm. After grinding, we maintained the food-waste batches at −20°C before using them in the bottle tests.

**Table 2.** Food-waste composition assignment of the sorted food-waste items. The food-waste items were categorized as either: carbohydrate-rich, lipid-rich, or protein-rich, depending on the component comprising the largest mass fraction (*w/w*). Two separate batches of protein-rich food waste were collected during the course of the large-media-bottle experiment.

| Carbohydrate-rich |  | Lipid-rich |  | Protein-rich I |  | Protein-rich II |  |
| --- | --- | --- | --- | --- | --- | --- | --- |
| [Item] | [% Mass] | [Item] | [% Mass] | [Item] | [% Mass] | [Item] | [% Mass] |
| Fruits | 15.4% | Olives | 14.1% | Red meat | 8.7% | Red meat | 8.8% |
| Vegetables | 47.1% | Fatty meats | 34.7% | Poultry | 8.0% | Poultry | 9.7% |
| Breads/Grains | 37.5% | Cheeses | 16.3% | Eggs | 40.0% | Eggs | 78.9% |
|  |  | Butter/Fats | 34.9% | Pork | 27.0% | Pork | 2.5% |
|  |  |  |  | Fish | 16.3% |  |  |

#### Operating conditions and sampling

For each experiment, we operated and sampled the bottles for an operating period of 15 days. To achieve the prescribed temperature condition, we kept the bottles in temperature-controlled, walk-in environmental chambers. The bottles were stored in the chambers for the duration of the operating period, except during biogas measurement and feeding, when the bottles were temporarily removed from the chambers. The biogas measurements, sampling, and feeding process, together, required approximately 10 min. The biogas measurements were taken *prior* to the feeding step. We measured the total biogas production volume on a daily basis *via* the gas-displacement method using a 30-mL glass syringe (Chemglass Life Sciences Co., Vineland, NJ). The resulting biogas volumes were then normalized to standard temperature and pressure (*i.e*., 273.15 K, 1.01 10^5^ Pa), assuming ideal gas conditions. In addition, we measured biogas composition (*i.e*., hydrogen [H_2_], methane [CH_4_], and carbon dioxide [CO_2_]) once per day for the first three days, and then every third day until the end of the operating period. We followed the same sampling schedule for most of the liquid-phase measurements, including: pH; ammonia; COD; and short-chain carboxylates (SCCs). However, in the case of the solids analysis, because it requires relatively large sample volumes, we only collected samples for these measurements on the first and last day of the experiment.

The amount of food waste fed during each feeding cycle also depended on the experimental feeding strategy (**Table 1**). For the ‘Batch’ feeding strategy for four experimental combinations, we filled the entire working volume (*i.e*., 3.75 L) of the bottle with food waste once at the very beginning of the incubation period (*i.e*., Day 0). For the two ‘Semi-Batch’ strategies (anoxic or semi-oxic), we fed 536 mL of food waste to the bottle on Days 0, 1, 2, 3, 6, 9, and 12, which resulted in an equivalent final working volume compared to the batch feeding strategy. For one of the semi-batch feeding strategies for four experimental combinations, which we refer to as ‘Semi-Batch: Semi-oxic’ (**Table 1**), we intentionally exposed the stored food waste to the atmosphere by opening the bottle lid during the feeding process. We then further allowed the headspace to equilibrate with the atmosphere for several minutes. Our intent was to determine the effect of periodic atmospheric O_2_ exposure. For the other semi-batch strategy for another four experimental combinations, which we refer to as ‘Semi-Batch: Anoxic’ (**Table 1**), we rigorously sparged the headspace during the feeding process to maintain anoxic conditions. Here, we intended to isolate the effect of a batch *vs*. semi-batch feeding regime by eliminating any confounding effects caused by O_2_ exposure.

### Biological-methane-potential experiments

At the end of the operating period of the storage experiment, we randomly selected one large-media bottle out of three possibilities (before knowing the fermentation outcome). Each of the 24 samples (12 before and 12 after) was tested in triplicate by the biological-methane-potential (BMP) method, resulting in 72 BMP-test bottles with a 250-mL volume. We followed the BMP method outlined in Labatut et al. ^5^, which is an adaptation of the original BMP method described by Owen et al. (1979). All of the materials and methods and the detailed results for these BMP experiments are described in the **Supporting Information**.

### Analytical methods

Biogas composition from the large-media-bottle storage experiment and the BMPs was measured using two gas chromatograph (GC) systems (Model: 8610C, SRI Instruments, Torrance, CA). Both GC systems were equipped with a packed column (0.3-m HaySep-D packed Teflon; Restek, Bellefonte, PA) and thermal conductivity detector and operated at isothermal conditions (40°C), however, one GC used He as carrier gas and was used to measure N_2_, CH_4_, and CO_2_, while the other GC used N_2_ as carrier gas and was used to measure H_2_. A third GC (6890 Agilent, Santa Clara, CA), which was equipped with a capillary column (NUKOL, Fused Silica Capillary Column, 15 m × 0.53 mm × 0.50 μm film thickness; Supelco Inc., Bellefonte, PA) and flame ionization detector, was used to measure the SCCs: acetate, propionate, and *n*-butyrate, using He as carrier gas according to Usack & Angenent (2015) ^6^. For lactate measurement, we used high performance liquid chromatography (HPLC) (Prominence, Shimadzu, Columbia, MD, USA). The HPLC was equipped with an RID-10A refractive index detector and Bio-Rad Aminex HPX-87H column operated at 65°C with 5 mM sulfuric acid as the mobile phase at a flow rate of 0.6 mL min^−1^. For all remaining analyses, we followed the methods outlined in ‘Standard Methods for the Examination of Water and Wastewater’ ^29^. Specifically, pH and ammonia concentration were measured using a pH meter (Model Orion 4-Star Plus; Thermo-Scientific, Waltham, MA) and ion-specific electrode (Model Orion 95-12; Thermo-Scientific, Waltham, MA), respectively. Also, we used the closed reflux titration method for COD analysis. In the case of soluble COD, we first centrifuged the samples for 10 min at 10,000 rpm, and then filtered the supernatant using a 0.2-μm Nylon filter (VWR, Radnor, PA).

## Results and discussion

### Food-waste and inoculum characteristics for the storage experiment

Preliminary work is shown in the **Supporting Information** (**Table S1-S5** and **Fig. S2-S6**), and this had informed us on the experimental handling of the bottles and also on which factors to include for the storage experiment. The ability to freeze food waste without affecting the outcome of fermentation meant that we could keep large batches of food waste for long time periods and use them across experiments. Furthermore, we used unmixed conditions in the storage experiment, reducing the complexity. In addition, the finding that food-waste composition had an effect on fermentation product spectra, apprised our decision to separate food waste into three bins: carbohydrate-rich, lipid-rich, or protein-rich food waste (**Table 2**), with the goal to study composition-related effects. Therefore, we used pre-frozen *and* sorted food waste as feedstock for the storage experiments. The initial pH values of the carbohydrate-rich, lipid-rich, and protein-rich food waste were mildly acidic to near neutral (5.7-6.6) (**Table 3**). The inoculum sample from a commercial-scale food-waste storage system had a lower pH value of 3.6 (**Table 3**). The relatively low ratios of SCOD TCOD^−1^ that were between 0.19-0.38 for carbohydrate-rich, lipid-rich, and protein-rich food waste indicates that most of the substrate was initially in the particulate form. In contrast, the ratio of SCOD TCOD^−1^ for the inoculum sample was relatively high at 0.82 (**Table 3**). Compared to the food waste, the inoculum sample contained a considerably higher concentration of carboxylates, such as acetate (3.4 g COD L^−1^), and especially lactate (58 g COD L^−1^) (**Table 3**). The lower pH value, higher ratio of SCOD TCOD^−1^, and higher concentrations of carboxylates for the inoculum sample compared to the food waste, indicated that the storage inoculum sample had undergone considerable hydrolysis and acidification (fermentation), while the food waste had not. The collections for two different protein-rich food-waste batches found no significant differences (p>0.53) in the fermentation product spectra (**Supporting Information** and **Fig. S7A-D**).

**Table 3.** Characterization of carbohydrate-rich, lipid-rich, protein-rich I, and protein-rich II food waste, and the inoculum. NOTE: All measured concentrations have been normalized to a common TS concentration of 100 g L^−1^ (10% *w/v*) (marked in bold font) to facilitate comparisons between food waste samples, because a 100 g L^−1^ TS concentration was used in the storage experiment. Errors represent the standard deviation. BD = below detection.

| Measured parameters | Carbohydrate-rich | Lipid-rich | Protein-rich I | Protein-rich II | Inoculum |
| --- | --- | --- | --- | --- | --- |
| pH* (*n*=1) | 6.05 | 5.67 | 6.43 | 6.55 | 3.60 |
| Ammonia (mg NH<sub>3</sub>-N L<sup>-1</sup>) (*n*=1) | 40 | 20 | 108 | 12 | 78 |
| VS (g L<sup>-1</sup>) (*n*=3) | 95 ± 2 | 92 ± 2 | 72 ± 1 | 32 ± 1 | 94 ± 22 |
| **TS (g L<sup>-1</sup>) (*n*=3)** | **100 ± 1** | **100 ± 1** | **100 ± 2** | **100 ± 8** | **100 ± 23** |
| VS TS<sup>-1</sup> (*n*=3) | 0.95 | 0.93 | 0.69 | 0.33 | 0.94 |
| SCOD (g L<sup>-1</sup>) (*n*=3) | 57 ± 4 | 9 ± 4 | 14 | 10 ± 1 | 77 ± 3 |
| TCOD (g L<sup>-1</sup>) (*n*=3) | 91 ± 3 | 57 ± 3 | 67 ± 4 | 38 ± 2 | 94 ± 11 |
| SCOD TCOD<sup>-1</sup> (*n*=3) | 0.38 | 0.19 | 0.21 | 0.26 | 0.82 |
| Acetate** (mg COD L<sup>-1</sup>) (*n*=1) | 1936 | 474 | 238 | 175 | 3338 |
| Propionate** (mg COD L<sup>-1</sup>) (*n*=1) | 137 | BD | 180 | BD | BD |
| Lactate** (mg COD L<sup>-1</sup>) (*n*=1) | 3527 | 572 | 1549 | 1882 | 58143 |
| *n*-Butyrate** (mg COD L<sup>-1</sup>) (*n*=1) | BD | 117 | 222 | 62 | 214 |

\* The pH was measured before dilution to a 100 g L<sup>-1</sup> TS concentration.
\*\* The carboxylates in this study refer to the combination of the undissociated and dissociated species.

### Food-waste composition, time, and temperature influenced each of the fermentation product spectra that we measured

Food-waste composition, time, and storage temperature significantly influenced every dependent variable that we tested in that order of importance (p<0.05 in **Fig. S8**). Indeed, PCA for the 288 sampling points showed a clear clustering pattern when the temperature (**Fig. S9A**) and food-waste composition (**Fig. S9B**) were highlighted. The strong effect of food-waste composition and time makes sense because carbohydrate-rich, lipid-rich, and protein-rich food waste have considerably different hydrolysis rates and degradability characteristics ^5,30,31^. The strong effect of the temperature is in agreement with the rates of degradation of complex substrates in anaerobic bioprocesses under different temperature regimes ^32^. In our study, the factor of feeding strategy affected most of the dependent variables, but three variables were not significantly affected by this factor (H_2_ production, gas production, and ammonia concentration in **Fig. S8**).

Even though inoculation is not necessary, we tested the factor of inoculation from a real food-waste storage facility to understand whether inoculation changes the fermentation outcome or whether it is kinetically controlled (*i.e*., whether the inoculum has an effect on fermentation time). We never measured any methane production during our two-week storage experiments, which is in agreement with ensilage research ^22^, and thus also not with inoculation. In addition, inoculation was the factor that had the least influence on the dependent variables compared to the other factors (**Fig. S8**). Importantly, the inoculation did not change the fermentation time. Still, the pH, ammonia concentration, lactate concentration, CO_2_ production, acetate concentration, and *n*-butyrate concentration were significantly affected by inoculum addition (**Fig. S8**). Because primarily the liquid concentrations were affected in the large-media bottles, we believe that the lower pH value, higher ammonia concentration, higher acetate concentration, and higher lactate concentration for the inoculum solution (chemical composition) changed the measured variables to a much greater degree than did the microbes present in the inoculum (microbial composition) (**Table 3**). This observation indicates again that inoculation of a microbiome before a two-week food storage period is not necessary, and is in agreement with several other studies ^13,19^. Our PCA for 288 sampling points agrees with the MANOVA results, because we did not observe a clear clustering when the feeding strategy (**Fig. S9C**) and inoculation (**Fig. S9D**) were highlighted. In regards to each of the dependent variables, we observed that the angle of the line and arrow for lactate concentration was considerably different from the other variables (**Fig. S9E**), indicating a unique effect of factors on lactate concentrations. On the other hand, the factors affected the individual fermentation gases and total gas production similarly, because they clustered together (**Fig. S9E**). In addition, acetate, propionate, *n*-butyrate, and ammonia concentrations clustered together as well (**Fig. S9E**).

The MANOVA test was also able to detect differences based on the interactions of multiple factors. Most of the two-way and three-way interactions (food-waste composition*feeding strategy, feeding strategy*time, food-waste composition*time, time*temperature, food-waste composition*feeding strategy*time, and time*inoculum) also significantly affected many dependent variables (**Fig. S8**). This suggests that the different factors influenced each other. This led us to develop interaction plots with two factors (temperature and time, feeding strategy and time, and inoculation and time) for each dependent variable and between the different food-waste compositions (**Fig. 1-2**, **Fig. S10-S11**, and **Fig. S12-S13**). These figures are all based on the *modeled* data, which represents the *measured* data well, but may not be identical (**Table S6)**.

**Figure 1.**
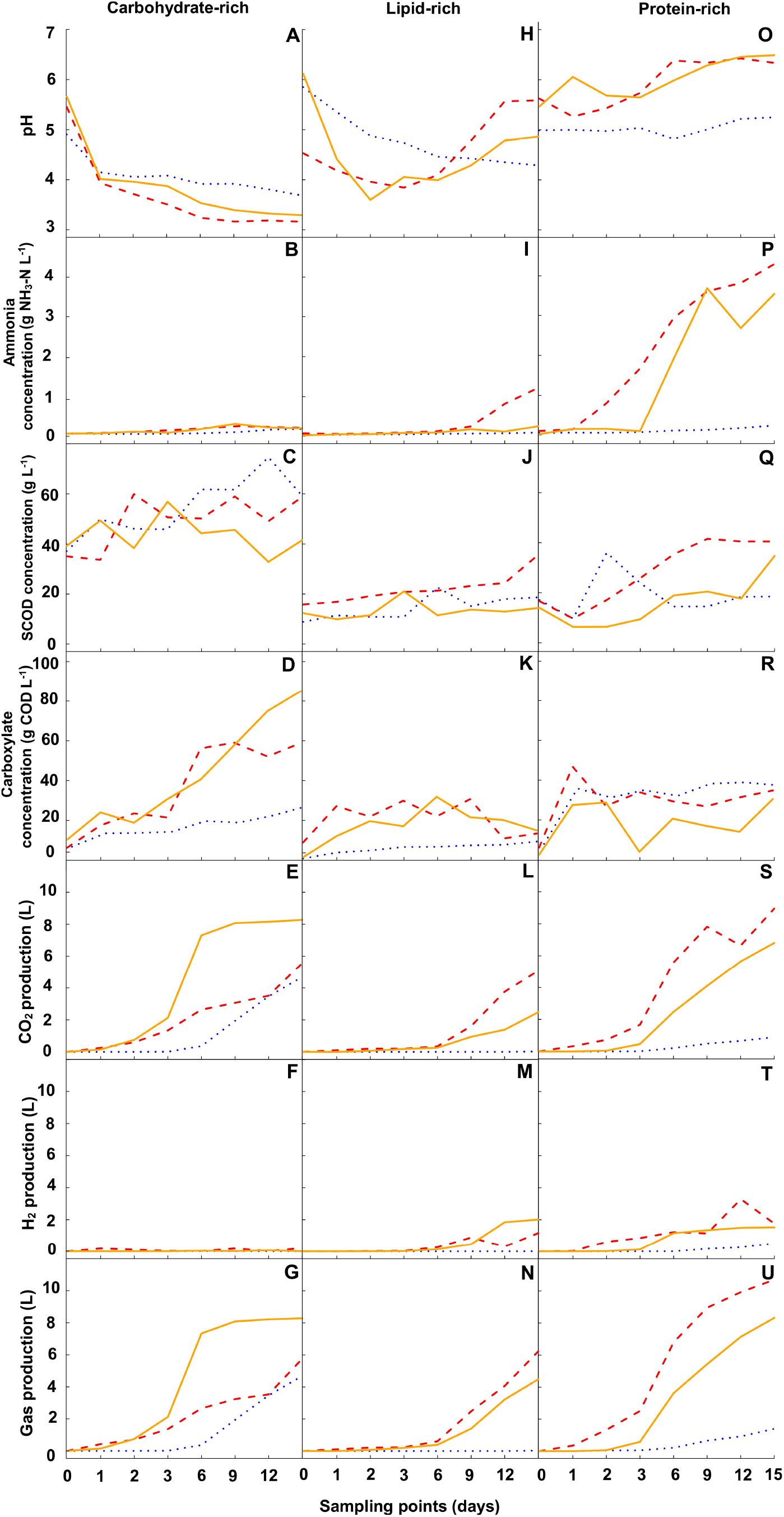
Twenty-one interaction plots for each dependent variable and for each of three food-waste compositions for eight sampling points throughout the operating period. Each of these plots is made with one of three different linear regression models (one for each food-waste composition) with the interacting factors temperature *plus* time. The resulting regression line with the response variables represents the predicted daily dependent variable throughout the operating period of the large-media-bottle-experiment for one temperature with a total of three lines for each temperature (blue dotted line = 15°C; orange solid line = 25°C; and red dashed line = 35°C). Here, we show seven different dependent variables (pH, ammonia concentration, SCOD concentration, total carboxylate concentration, CO_2_ production, H_2_ production, and total gas production) throughout the operating time for three different types of food waste (carbohydrate-rich, lipid-rich, and protein-rich), resulting in 21 panels (**A-U**). Curves that are not parallel within one panel indicate interaction between temperature and time. Overall, changes in temperature do not have a linear effect on the dependent variable. Note that the x-axis (time) is intentionally not to scale to identify that this is data based on a model.

**Figure 2.**
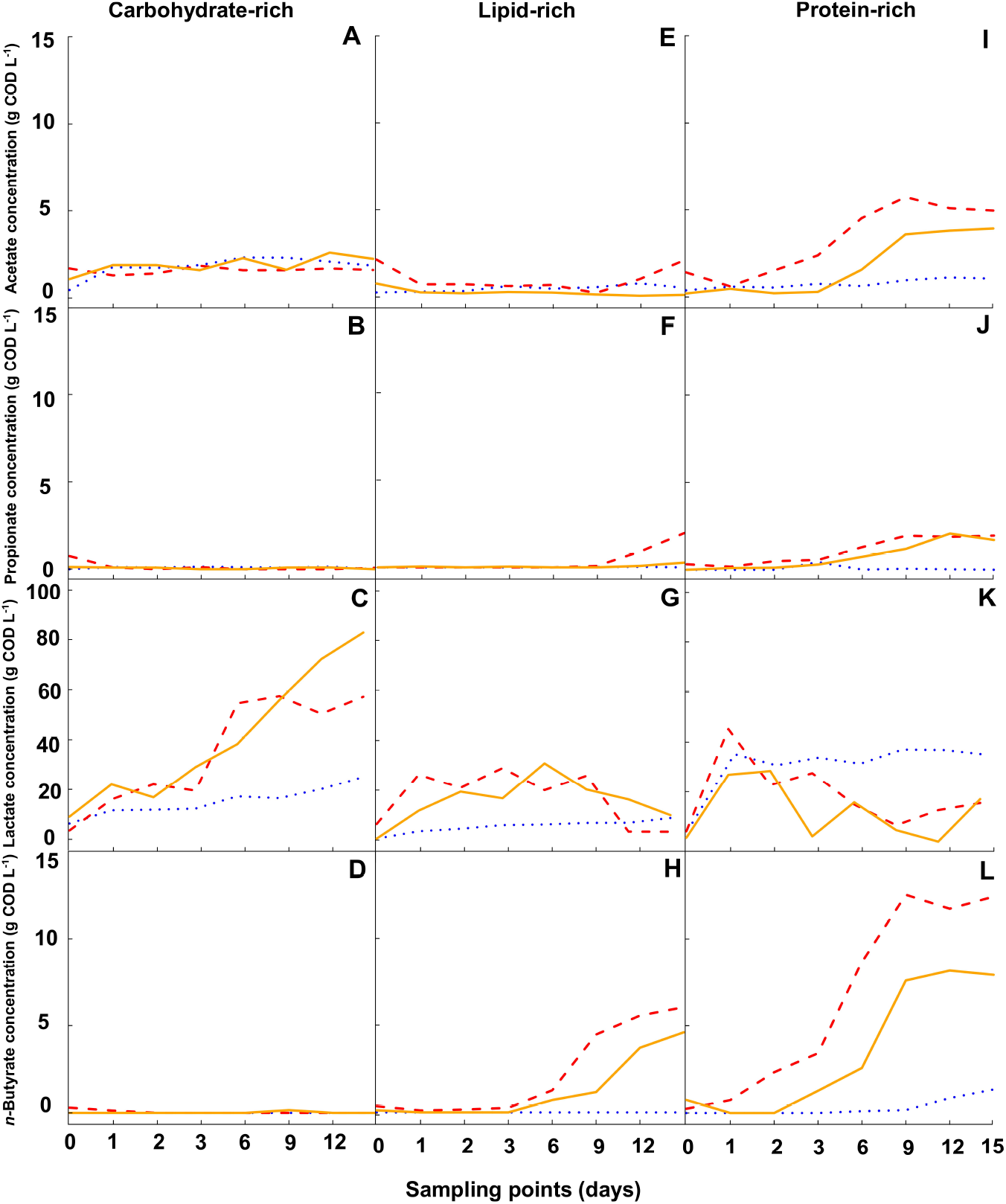
Twelve interaction plots for each of the SCC concentrations (dependent variables) throughout the operating period. The same linear regression models with the interacting factors temperature *plus* time were used as in Fig. 1. Three lines for each temperature (blue dotted line = 15°C; orange solid line = 25°C; and red dashed line = 35°C) in each panel show the predicted SCC concentrations for acetate, *n*-propionate, lactate, and *n*-butyrate throughout the operating period for each temperature. With three different types of food waste (carbohydrate-rich, lipid-rich, and protein-rich), this resulted in 12 panels (**A-L**). Curves that are not parallel within one panel indicate interaction between temperature and time. Overall, changes in temperature do not have a linear effect on the SCC concentrations. Note that the x-axis (time) is intentionally not to scale to identify that this is data based on a model.

### Effects of food-waste composition and temperature on fermentation product spectra

We focused on temperature and food-waste composition, because these factors *plus* time affected all of the dependent variables (*i.e*., the fermentation product spectra), which was not the case for feeding strategy and inoculation (**Fig. S8-S9**). Therefore, the results for the factors feed strategy and food-waste composition *plus* time and inoculation and food-waste composition *plus* time are shown in the **Supporting Information** (**Fig. S10-S13**). We discuss the fermentation product spectra based on interaction plots with the factor temperature *plus* time for the food-waste compositions to show the relationship between temperature and time on the dependent variables for each food-waste composition. These plots were made by fitting data using linear regression models (**Eq. 1**), and including: 1) eight sampling points for each of the dependent variables; and 2) the data from each of the food-waste compositions separately. By doing so, we obtained three regression lines (one for each temperature) in each panel with specific response variables (*e.g*., pH in **Fig. 1A**). Both the temperature and time were required to predict the dependent variables because food-waste degradation is complex, and temperature alone does not linearly affect the fermentation product spectra. Linear regression connected the measured sampling points (Days 0, 1, 2, 3, 6, 9, 12, and 15) with the non-measured points (Days 4, 5, 7, 8, 10, 11, 13, 14). To visualize the model data, we first used seven dependent variables for the three different food-waste compositions, resulting in 21 panels (**Fig. 1A-U**). Next, we plotted 12 panels (**Fig. 2A-L**) for the four different carboxylate concentrations as dependent variables.

#### Rapid acidification of carbohydrate-rich food waste

In general, carbohydrate-rich food waste experienced rapid acidification during the first three days of storage, regardless of temperature (**Fig. 1A**). This rapid acidification was caused primarily by the production of carboxylates by homofermentation (**Fig. 1D**), because the fermentation product consisted primarily of lactate (>90%) and only minor amounts of acetate (**Fig. 2A-D**). We did not observe saccharolytic clostridial fermentation to *n*-butyrate as a secondary fermentation during these days (**Fig. 2D**). The lower pH values reduced the subsequent acidification rate, however, the decreases in pH values continued until the end of the operating period for all temperatures, while the gas production also continued (**Fig. 1A-G**). Regardless, the carbohydrate-rich food waste became quickly too acidic (<4.5) for heterofermentation and/or saccharolytic clostridia fermentation of lactate into other fermentation products (*i.e*., secondary fermentation), which also prevented the production of H_2_ gas (**Fig. 1F**). The lowest pH was achieved at 35°C (**Fig. 1A**) and reached a value of 3.2 at the end of the operating period (Comb. 9 in **Table S6**). However, the maximum concentration of carboxylates and total gas production were achieved at 25°C (**Fig. 1D-G**), identifying the non-linear behavior of temperature. Due to the rapid acidification of carbohydrate-rich food waste *via* mostly homofermentation, the efficient conversion of this type of food waste during storage is halted in a similar fashion as ensilage of plant materials – through the rapid production of lactate ^33^, even preventing the possible conversion of lactate ^22^.

#### Rapid acidification of lipid-rich food waste was followed by a pH recovery

In contrast to carbohydrate-rich food waste, the rate of fermentation for lipid-rich food waste increased with increasing temperatures (**Fig. 1K-L,N**). This can partly be explained by the solubilization of the lipids into solution, which, when rate limiting, is generally accepted to occur faster at higher temperatures. Solubilization is followed by hydrolysis of triglycerides into glycerol and long-chain fatty acids (LCFAs). Therefore, the rate limitation of hydrolysis could also explain the higher rate of fermentation with higher temperatures. The relatively quick pH drop to 4.0 and 3.8 on Day 3 for the 25°C and 35°C experimental conditions, respectively (**Fig. 1H**), can be attributed to the relative fast fermentation rate for glycerol (a monosaccharide) to produce carboxylates (**Fig. 1K**), consisting mostly of lactate (**Fig. 2E-H**). In contrast to carbohydrate-rich food waste, however, the pH value did not drop as much, and even increased again for the 25°C and 35°C experimental conditions at the end of the operating period (**Fig. 1H**), with a final pH level of 5.6 at 35°C (Comb. 10 in **Table S6**). The pH did not drop enough to prevent saccharolytic clostridial fermentation of 2 moles of lactate into 1 mole of *n*-butyrate, 2 moles of CO_2_, and 2 moles of H_2_ ^16,17^ (**Fig. 2G-H** and **Fig. 1L-M**). This conversion explains the increase in pH, because: 1) 2 moles of lactate are converted to 1 mole of *n*-butyrate; and 2) lactate is a stronger acid with a lower pKa than *n*-butyrate ^22^. At the coldest temperature of 15°C, the pH decreased at a considerably slower rate compared to the 25°C and 35°C conditions (**Fig. 1H**), but the other fermentation pathways were not initiated within the operating period (**Fig. 1I-N**).

The higher pH for the 25°C and 35°C experimental conditions had secondary fermentation made possible. Besides *n*-butyrate and H_2_, however, some propionate was also produced after the increase in the pH level (**Fig. 1H,M** and **Fig. 2F,H**), which had not occurred noticeably during carbohydrate-rich food waste storage (**Fig. 1K** and **Fig. 2B,D**). For both the 25°C and 35°C experimental conditions, we also observed that the lactate concentration (and thus the total carboxylate concentration) first increased and then decreased after Day 6 (**Fig. 1K** and **Fig. 4G**), which coincided with the production of CO_2_, H_2_, propionate, and *n*-butyrate (**Fig. 1L-M** and **Fig. 2F, H**). The decrease in lactate concentration was considerably higher than the increase in *n*-butyrate concentration (note the different scales in **Fig. 2G-H**), and therefore additional microbial pathways must have become active at the mildly acidic pH levels of ∼5.7 to metabolize the lactate (**Fig. 1H**). We speculate that besides saccharolytic clostridial fermentation, the acrylate pathway to convert lactate into propionate may have become active ^34–36^, explaining the production of propionate (**Fig. 2F**). However, more work would be necessary to ascertain which microbial pathways were responsible.

#### Production of ammonia during protein-rich food-waste storage sustained near-neutral pH values

During the first two days of the operating period, lactate production during the protein-rich food-waste storage was rapid (**Fig. 2K**), which was similar for all food-waste compositions (**Fig. 2C, G**). For only the protein-rich food waste, however, the temperature did not affect the lactate production rate during the initial couple of days (**Fig. 2C,K,G**). Even though lactate production was rapid during the protein-rich food-waste storage, the pH did not drop substantially during the first two days of the operating period (**Fig. 1O**). The storage of protein-rich food waste was characterized by a considerable increase in the ammonia concentration at the warmer temperatures of 25°C and 35°C (**Fig. 1P**), which increased the pH levels to ∼6.5 at the end of the operating period (**Fig. 1O**). This increase in the pH level can be explained by the production of ammonia from proteolytic clostridial fermentation of, for example, lysine to acetic acid, *n*-butyric acid, and ammonia at pH levels that are higher than 4.5 ^16,17^ (**Fig. 1P** and **Fig. 2I,L**). Specifically, ammonia combines with water and CO_2_ to form ammonia bicarbonate, which is a pH buffer ^4^ The maximum achieved ammonia concentration was 4.7 g NH_3_-N L^−1^ at 35°C (Comb. 12 in **Table S6**). The pH levels had already increased to close to 6 after Day 3 (**Fig. 1O**), which made the same fermentation pathways possible for the 25°C and 35°C experimental conditions as we observed for the lipid-rich food-waste storage. We observed increases in the acetate, propionate, and *n*-butyrate concentrations and the cumulative CO_2_ and H_2_ (gas) production (**Fig. 1S-U** and **Fig. 2I-L**), while the lactate concentration decreased again (**Fig. 2K**). Due to the highest pH levels in the protein-rich food-waste storage, the highest cumulative H_2_ and *n*-butyrate production of 3 L and 13 g COD L^−1^, respectively, was achieved at 35°C (**Fig. 1T** and **Fig. 2L**). The temperature did not have a linear effect on the production of these fermentation products. At the lowest temperature of 15°C, we did not observe an increase in ammonia concentration (**Fig. 1P**). As a result, the pH level remained fairly steady between 5.0-5.3 (**Fig. 1O**). Lactate production leveled off almost immediately from the start (**Fig. 2K**).

### Important fermentation products for chemical and/or energy production

#### Acidic conditions promoted lactate production

Lactate is a common platform chemical, which has been used for years mainly as a preservative in the food industry. It also has other applications in the textile, cosmetic, and pharmaceutical industries ^37^. Recently, there has been an increased attention for lactate as: 1) a feedstock for bio-plastics ^38^; and 2) as an electron donor for microbial chain elongation with reverse β-oxidation to produce medium-chain carboxylates (caproate [*n*-hexanoate] and caprylate [*n*-octanoate]) that are used as feed, food, antimicrobials, and lubricants among other uses ^36^. Lactate was the dominant carboxylate produced for *each* of the food-waste compositions, and was produced relatively quickly during the operating period (**Fig. 2**). Still, the factor food-waste composition had a large effect on lactate production for the storage experiment, while the other factors *plus* time also affected lactate production (**Eq. S1**).

The highest lactate production in our storage experiment occurred for the carbohydrate-rich conditions at 25°C and 35°C (**Fig. 2C,G,K**). Specifically, we achieved a maximum average lactate concentration of 83 g COD L^−1^ (77 g L^−1^ or ∼7.7% *w/v*), which represented 97% of the total carboxylate concentration (**Fig. 2A-D**), at 25°C and a pH of 3.3 with carbohydrate-rich food waste in a large-serum bottle that was operated as a batch with inoculum (Comb. 5 in **Table S6**). For this combination, the maximum average lactate COD selectivity was 90% (based on the total COD that was originally in the food waste), which is similar to the maximum selectivity for lactate in another batch of a food-waste fermentation study ^39^. Our relatively high selectivity was achieved without product removal ^40^, but with substrate dilution ^41^ by water addition in the grinding system. The lactate became the main electron sink without the presence of methanogenesis ^42^. Subsequently, the quick acidification by lactate inactivated the microbiome (pickling or conservation at a low pH level of 3-4), resulting in the absence of H_2_ gas or other intermediates such as *n*-butyrate. Indeed, two other studies involving food waste found that when the pH declined to 4.5-5.0, H_2_ production was inhibited while lactate was produced ^43,44^

This subsequent lactate conversion into other fermentation products through saccharolytic clostridial fermentation and *via* the acrylate pathway did occur at the higher pH levels for lipid-rich and protein-rich food waste. For example, we observed the production of acetate, propionate, *n*-butyrate, and H_2_ from lactate for protein-rich food-waste storage at a pH level of 6-7 for the 25°C and 35°C experimental conditions (**Fig. 1O**, **T** and **Fig. 2I-L**). This lactate conversion process was considerably slower at the lower temperature of 15°C compared to 25°C and 35°C (**Fig. 1L**), and thus a lower temperature can prevent lactate conversion. Another study with auto-inoculation of food waste at a temperature of 37°C also found that lactate was only further converted into SCCs at a higher pH level of 6.0 and not at pH level of 4.0 and 5.0 ^45^. For a biorefinery concept aimed to produce lactate, a pre-separation step of the bulk food waste into a carbohydrate-rich food-waste bin maybe advisable. By using primarily carbohydrate-rich food waste, quick acidification due to lactate production (even at the lower temperatures) without a likely return to more basic conditions, would prevent the subsequent lactate conversion during uncontrolled storage, which is an advantageous situation during food storage. Another possibility is to maintain a pH below 4.5 by installing pH control for the storage tank,

#### A low temperature or acidic condition curtailed H_2_ production

H_2_ gas production can be regarded as a negative during food-waste storage for several reasons. First, H_2_ is an ignitable gas at partial pressures greater than 5%, which could pose hazards to a decentralized storage system. Second, large volumes of H_2_ gas production could lead to a COD loss from the food-waste slurry, which reduces the methane potential of food waste ^16^. The factor food-waste composition played a large role for H_2_ gas production. For example, H_2_ production was not noticeable for the carbohydrate-rich conditions, while H_2_ production occurred in the lipid-rich conditions at warmer temperatures (25°C, 35°C) (**Fig. 1M**), and increased with increasing temperature in protein-rich conditions (**Fig. 1T**), with the factors temperature and food-waste composition describing the relationship (**Eq. S2**). The H_2_ production was related to the pH level during the experiments for the same reasons as for lactate conversion. (**Fig. 1H,O,M,T**). Other studies had already shown that H_2_ production in anaerobic environments was gradually inhibited below the optimum pH range of 5.0-6.0 ^46^.

Even at the highest average cumulative H_2_ gas production of 1.95 L at a final average pH value of 6.2 during 15 days at 35°C, with protein-rich food waste for a batch feeding strategy and with inoculation (Comb. 11 in **Table S6**), only a H_2_ COD selectivity of ∼1% was achieved, which is relatively low. This explains why we did not measure a significant loss of TCOD concentration in large-media bottles (**Table S6**). However, the maximum average H_2_ concentration that we achieved in the headspace of a large-media bottle was 51.4% with lipid-rich food waste that was fed as a batch and without inoculation at 25°C (Comb. 6 in **Table S6**), which is higher than the ignitable limit. We observed that H_2_ production for each food-waste composition was largely prevented at a low temperature of 15°C. To prevent hydrogen formation, however, cooling a large volume of food slurry may not be economical during warm weather periods or when the slurry is stored indoors. Another prevention method is to maintain a relatively low pH value below 4.5. This was intrinsically accomplished by using only carbohydrate-rich food waste, but installing pH control for the storage tank would be another method to prevent hydrogen production.

### Methane selectivity increased for food-waste storage at high lactate concentrations

We conducted BMP assays for each of the 12 experimental combinations, before and after storage, to determine whether the specific storage conditions affected the anaerobic methane selectivity (*i.e*., volume of methane production *per* mass of TCOD added) for the food waste (details about the methods and results in the **Supporting Information**). We found that in the three combinations with a significant increase in the methane selectivity (**Fig. S14**), the concentrations of lactate were also high at the end of the operating period (Comb. 5, 9, and 12 in **Table S6**). We observed a significant lower methane selectivity for only one combination (Comb. 2; 15°C with carbohydrate-rich food waste in **Fig. S14**), but we do not know the reason.

## Supporting Information

The Supporting Information is available free of charge at xxx. It contains: 30 pages, 10 sections, 14 figures, 6 tables, and 2 equations.

## Synopsis

The decentralized grinding, diluting, and storage of food waste is promising as a pretreatment step to recover the carbon.

## Author Contributions

LTA and MPL conceived the project and LTA, JGU, and SED designed the study. SED performed the lab experiments. JGB advised on the statistical analyses. SED and LAH analyzed the data. SED and JGU prepared the figures and tables. SED, JGU, and LTA drafted the manuscript. LTA and MPK provided guidance. All authors edited the manuscript and approved the final manuscript.

## Notes

The authors declare no competing financial interest.

## Abbreviations

AD: anaerobic digestion
MANOVA: multivariate analysis of variance
BMP: biochemical methane potential
COD: chemical oxygen demand
GHG: greenhouse gas
LCFA: long chain fatty acid
MFFA: mixed-level fractional factorial analysis
SCCs: short-chain carboxylates
SCOD: soluble chemical oxygen demand
TCOD: total chemical oxygen demand
TS: total solids
VFA: volatile fatty acid
VS: volatile solids.

## Acknowledgements

This work was supported by a gift from InSinkErator, which is a subsidiary of Emerson, as part of their Grind2Energy^™^ food-waste-recycling program. The food-waste grinder and the inoculum used in this study were also gifts from InSinkErator. In addition, we thank the staff at Cornell Dining for providing food waste and Françoise Vermeylen (Cornell Statistical Consulting Unit) for statistical consulting (both at Cornell University, Ithaca, NY).

## References

1. U.S. EPA (2016) Advancing sustainable materials management: 2014 fact sheet assessing trends in material generation, recycling, composting, combustion with energy recovery and landfilling in the United States (United States Environmental Protection Agency, Washington, DC 20460), (Office of Land and Emergency Management (5306P)).

2. U.S. EPA (2016) Inventory of u.S. Greenhouse gas emissions and sinks: 1990 – 2014 (United States Environmental Protection Agency, Washington, DC 20460).

3. Cuéllar, A.D., and Webber, M.E. (2010). Wasted food, wasted energy: The embedded energy in food waste in the United States. Environ. Sci. Technol. 44, 6464–6469. https://doi.org/10.1021/es100310d.

4. Zhang, B., Zhang, L.L., Zhang, S.C., Shi, H.Z., and Cai, W.M. (2005). The influence of pH on hydrolysis and acidogenesis of kitchen wastes in two-phase anaerobic digestion. Environmental Technology 26, 329–339. https://doi.org/10.1080/09593332608618563.

5. Labatut, R.A., Angenent, L.T., and Scott, N.R. (2011). Biochemical methane potential and biodegradability of complex organic substrates. Bioresour. Technol. 102, 2255–2264. https://doi.org/10.1016/j.biortech.2010.10.035.

6. Usack, J.G., and Angenent, L.T. (2015). Comparing the inhibitory thresholds of dairy manure co-digesters after prolonged acclimation periods: Part 1 - performance and operating limits. Water Res. 87, 446–457. http://dx.doi.org/10.1016/j.watres.2015.05.055.

7. Usack, J.G., Gerber Van Doren, L., Posmanik, R., Labatut, R.A., Tester, J.W., and Angenent, L.T. (2018). An evaluation of anaerobic co-digestion implementation on new york state dairy farms using an environmental and economic life-cycle framework. Appl. Energy 211, 28–40. https://doi.org/10.1016/j.apenergy.2017.11.032.

8. ten Brummeler, E. (2000). Full scale experience with the biocel process. Water Sci. technol. 41, 299–304.

9. Mata-Alvarez, J., Mac, S., and Llabr, P. (2000). Anaerobic digestion of organic solid wastes. An overview of research achievements and perspectives. Biores. technol. 74.

10. Breure, A.M., Mooijman, K.A., and Andel, J.G.V. (1986). Protein degradation in anaerobic digestion: Influence of volatile fatty acids and carbohydrates on hydrolysis and acidogenic fermentation of gelatin. Appl. Microbiol. Biotechnol. 24, 426–431.

11. De Baere, L. (2000). Anaerobic digestion of solid waste: State-of-the-art. Water Sci. Technol. 41, 283–290.

12. Kim, D.-H., Wu, J., Jeong, K.-W., Kim, M.-S., and Shin, H.-S. (2011). Natural inducement of hydrogen from food waste by temperature control. Int. J. Hydrogen Energy 36, 10666–10673. https://doi.org/10.1016/j.ijhydene.2011.05.153.

13. Kim, D.-H., Kim, S.-H., and Shin, H.-S. (2009). Hydrogen fermentation of food waste without inoculum addition. Enzyme Microb. Technol. 45, 181–187. https://doi.org/10.1016/j.enzmictec.2009.06.013.

14. Kim, D.-H., Kim, S.-H., Jung, K.-W., Kim, M.-S., and Shin, H.-S. (2011). Effect of initial pH independent of operational pH on hydrogen fermentation of food waste. Bioresour. Technol. 102, 8646–8652. https://doi.org/10.1016/j.biortech.2011.03.030.

15. Kandler, O. (1983). Carbohydrate metabolism in lactic acid bacteria. Antonie Van Leeuwenhoek 49, 209–224. https://doi.org/10.1007/BF00399499.

16. Teixeira Franco, R., Buffière, P., and Bayard, R. (2016). Ensiling for biogas production: Critical parameters. A review. Biomass Bioenerg. 94, 94–104. https://doi.org/10.1016/j.biombioe.2016.08.014.

17. McDonald, P., Henderson, A.R., and Heron, S.J.E. (1991) The biochemistry of silage Chalcombe Publications. 340 pp.

18. Kim, D.-H., and Kim, M.-S. (2013). Development of a novel three-stage fermentation system converting food waste to hydrogen and methane. Bioresour. Technol. 127, 267–274. https://doi.org/10.1016/j.biortech.2012.09.088.

19. Kim, M.-S., Na, J.-G., Lee, M.-K., Ryu, H., Chang, Y.-K., Triolo, J.M., Yun, Y.-M., and Kim, D.-H. (2016). More value from food waste: Lactic acid and biogas recovery. Water Res. 96, 208–216. https://doi.org/10.1016/j.watres.2016.03.064.

20. Peinemann, J.C., Rhee, C., Shin, S.G., and Pleissner, D. (2020). Non-sterile fermentation of food waste with indigenous consortium and yeast – effects on microbial community and product spectrum. Bioresour. Technol. 306, 123175. https://doi.org/10.1016/j.biortech.2020.123175.

21. Daeschel, M.A., Andersson, R.E., and Fleming, H.P. (1987). Microbial ecology of fermenting plant materials: Presented at the second symposium on lactic acid bacteria-genetics, metabolism and applications, 22-25 September 1987, Wageningen, the Netherlands. FEMS Microbiol. Rev. 3, 357–367. https://doi.org/10.1111/j.1574-6968.1987.tb02472.x.

22. Hillion, M.-L., Moscoviz, R., Trably, E., Leblanc, Y., Bernet, N., Torrijos, M., and Escudié, R. (2018). Co-ensiling as a new technique for long-term storage of agro-industrial waste with low sugar content prior to anaerobic digestion. Waste Manag. 71, 147–155. https://doi.org/10.1016/j.wasman.2017.10.024.

23. Ambye-Jensen, M., Johansen, K.S., Didion, T., Kádár, Z., Schmidt, J.E., and Meyer, A.S. (2013). Ensiling as biological pretreatment of grass (*Festulolium hykor*): The effect of composition, dry matter, and inocula on cellulose convertibility. Biomass Bioenerg. 58, 303–312. https://doi.org/10.1016/j.biombioe.2013.08.015.

24. Jawad, A.H., Alkarkhi, A.F.M., Jason, O.C., Easa, A.M., and Nik Norulaini, N.A. (2013). Production of the lactic acid from mango peel waste – factorial experiment. J. King Saud Univ. Sci. 25, 39–45. https://doi.org/10.1016/j.jksus.2012.04.001.

25. Wang, S., Huang, G.H., and Zhou, Y. (2016). A fractional-factorial probabilistic-possibilistic optimization framework for planning water resources management systems with multi-level parametric interactions. J. Environ. Manag. 172, 97–106. https://doi.org/10.1016/j.jenvman.2016.02.019.

26. Dauneau, P., and Perez-martinez, G. (1997). Fractional factorial design and multiple linear regression to optimise extraction of volatiles from a lactobacillus plantarum bacterial suspension using purge and trap. J. Chromat. A 775, 225–230.

27. Kutner, M.H., Nachtsheim, C.J., Neter, J., and Li, W. (2015) Applied linear regression models MacGraw-Hill Irwin Applied linear regression models 5 Ed.

28. USDA (2015) USDA national nutrient database for standard reference, release 27 (Agricultural Research Service, Washington, DC), (Nutrient data laboratory).

29. APHA (1999) Standard methods for the examination of water and wastewater American Public Health Association; American Water Works Association; Water Environment Federation Standard methods for the examination of water and wastewater 20th Ed p 5.

30. Neves, L., Gonçalo, E., Oliveira, R., and Alves, M.M. (2008). Influence of composition on the biomethanation potential of restaurant waste at mesophilic temperatures. Waste Manag. 28, 965–972. https://doi.org/10.1016/j.wasman.2007.03.031.

31. Christ, O., Wilderer, P.A., Angerhöfer, R., and Faulstich, M. (2000). Mathematical modeling of the hydrolysis of anaerobic processes. Water Sci. Technol. 41, 61–65.

32. Pavlostathis, S.G., and Giraldo-Gomez, E. (1991). Kinetics of anaerobic treatment: A critical review. Crit. Rev. Environ. Contr. 21, 411–490. https://doi.org/10.1080/10643389109388424.

33. Driehuis, F., Oude Elferink, S.J.W.H., and Spoelstra, S.F. (1999). Anaerobic lactic acid degradation during ensilage of whole crop maize inoculated with lactobacillus buchneri inhibits yeast growth and improves aerobic stability. J. Appl. Microbiol. 87, 583–594.

34. Prins, R.A., and Meer, P. (1976). On the contribution of the acrylate pathway to the formation of propionate from lactate in the rumen of cattle. Antonie Van Leeuwenhoek 42, 25–31.

35. Prabhu, R., Altman, E., and Eiteman, M.A. (2012). Lactate and acrylate metabolism by *Megasphaera elsdenii* under batch and steady-state conditions. Appl. Environ. Microbiol. 78, 8564–8570.

36. Kucek, L.A., Nguyen, M., and Angenent, L.T. (2016). Conversion of L-lactate into *n*-caproate by a continuously fed reactor microbiome. Water Res. 93, 163–171. http://dx.doi.org/10.1016/j.watres.2016.02.018.

37. Zhao, L., Liu, D.X., and Wang (2007). Effect of several factors on peracetic acid pretreatment of sugarcane bagasse for enzymatic hydrolysis. J. Chem. Technol. Biotechnol. 82, https://doi.org/1115-1121.10.1002/jctb.

38. Gao, C., Ma, C., and Xu, P. (2011). Biotechnological routes based on lactic acid production from biomass. Biotechnol. Adv. 29, 930–939. https://doi.org/10.1016/j.biotechadv.2011.07.022.

39. Peinemann, J.C., Demichelis, F., Fiore, S., and Pleissner, D. (2019). Techno-economic assessment of non-sterile batch and continuous production of lactic acid from food waste. Bioresour. Technol. 289, 121631. https://doi.org/10.1016/j.biortech.2019.121631.

40. Wang, X., Wang, Y., Zhang, X., Feng, H., and Xu, T. (2013). *In-situ* combination of fermentation and electrodialysis with bipolar membranes for the production of lactic acid: Continuous operation. Bioresour. Technol. 147, 442–448. https://doi.org/10.1016/j.biortech.2013.08.045.

41. Xu, J., Hao, J., Guzman, J.J.L., Spirito, C.M., Harroff, L.A., and Angenent, L.T. (2018). Temperature-phased conversion of acid whey waste into medium-chain carboxylic acids via lactic acid: No external e-donor. Joule 2, 280–295. https://doi.org/10.1016/j.joule.2017.11.008.

42. Fukuzaki, S., Chang, Y.J., Nishio, N., and Nagai, S. (1991). Characteristics of granular methanogenic sludge grown on lactate in a UASB reactor. J. Ferment. Bioeng. 72, 465–472.

43. Marone, A., Izzo, G., Mentuccia, L., Massini, G., Paganin, P., Rosa, S., Varrone, C., and Signorini, A. (2014). Vegetable waste as substrate and source of suitable microflora for bio-hydrogen production. Ren. Energy 68, 6–13. https://doi.org/10.1016/j.renene.2014.01.013.

44. Wang, X., and Zhao, Y. (2009). A bench scale study of fermentative hydrogen and methane production from food waste in integrated two-stage process. Int. J. Hydr. Energ. 34, 245–254. https://doi.org/10.1016/j.ijhydene.2008.09.100.

45. Tang, J., Wang, X.C., Hu, Y., Zhang, Y., and Li, Y. (2017). Effect of pH on lactic acid production from acidogenic fermentation of food waste with different types of inocula. Bioresour. Technol. 224, 544–552. https://doi.org/10.1016/j.biortech.2016.11.111.

46. Kapdan, I.K., and Kargi, F. (2006). Bio-hydrogen production from waste materials. Enzyme Microb. Technol. 38, 569–582. http://dx.doi.org/10.1016/j.enzmictec.2005.09.015.

