## Supporting Information for "Food waste as a resource: grinding, dilution, and storage as a pretreatment strategy to produce fermentation intermediates"

**This SI contains: 30 pages, 10 sections, 14 figures, 6 tables, and 2 equations**

### I) Large-media bottles for the storage experiment

We performed the storage experiment using 5-L laboratory borosilicate-glass large-media bottles (Duran, Mainz, Germany) with a final working volume of 3.75 L. A glassblower (Cornell University) added three 2.54-cm diameter sampling ports with outside glass threading to each bottle to facilitate: 1) liquid sampling; 2) headspace gas sampling; and 3) headspace sparging with nitrogen gas (**Fig. S1**). Finally, we made sure that each port was sealed using rubber septa and polytetrafluoroethylene (PTFE) liners, which we fixed tightly in place with threated plastic caps.

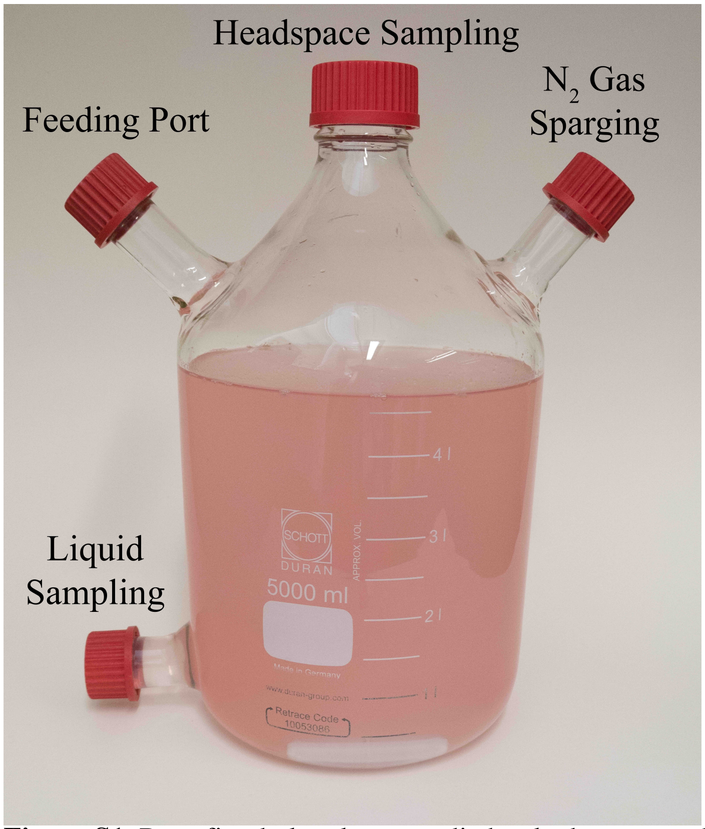


**Figure S1.** Retrofitted glass large-media bottle that we used for the storage experiments (not for the preliminary storage experiment with small-media bottles). The colored solution was added to the bottle to create contrast.

### II) Preliminary-storage experiments with small-media bottles

In **Section S1.1**, we provide a description of the general methodology that was applied for each preliminary experiment, while in **Sections S1.2-S1.6**, we explain in more detail the specific ways a given preliminary experiment deviated from this general method. In addition, in each section we present the relevant data and highlight the main statistical results from each experimental parameter for these preliminary experiments. For the preliminary-storage experiments we varied the following five experimental parameters: 1) storage temperature; 2) feeding strategy; 3) inoculation; 4) mixing; and 5) freezing. Note: the data pertaining to food-waste composition, which is one of the experimental parameters used for the large-bottle experiment is not presented here because no preliminary experiments were directly performed with different food-waste composition conditions. However, we were able to infer that food waste composition would have a significant effect on the fermentation process by comparing the results between the preliminary experiments presented. Each preliminary experiment involved a different batch of food waste with varied composition. An important observation of the preliminary-storage experiments, therefore, was to vary and control the food-waste composition as an additional experimental parameter. Another important finding from the preliminary experiments was that mixing and freezing did not have a statistically significant effect on the fermentation product spectrum during food storage.

#### Section S1.1: General approach for the preliminary-storage experiments

The food waste that we used for the preliminary-storage experiments was collected from the same dining hall at Cornell University as for the storage experiments. Immediately after collection, we ground the food waste with the InSinkerator® food disposal system (Model MSLV-7, Emerson Electric Co, Racine, WI) and then characterized its composition. The characterization measurements included: ammonia, total solids (TS), volatile solids (VS), pH, soluble chemical oxygen demand (SCOD), total COD (TCOD), acetate, propionate, and *n*-butyrate. Initially, we collected individual batches of food waste *on demand,* which we would store for short periods (~1-3 days) in a refrigerator (4°C) until use. This was performed because we had not yet determined the effects of freezing on food-waste storage. The concentrations for the SCCs were converted to mg COD $L^{-1}$ by multiplying the mM concentration (determined by GC or HPLC) with the molar mass and its theoretical oxygen demand (1.08 mg COD mg $acetate^{-1}$; 1.53 mg COD mg $propionate^{-1}$; 1.08 mg COD mg $lactate^{-1}$; and 1.84 mg COD mg *n*-$butyrate^{-1}$).

We used 150-mL glass media bottles or 250-mL glass media bottles with a working volume of 110 or 190 mL, respectively, for these preliminary-storage experiments. Each bottle was fed ground food waste diluted to 10% TS at the beginning of the incubation period and then the bottles were sealed. The duration of the incubation period varied depending on the time needed to achieve a stable system. Due to the smaller volume of these bottles, *in-situ* sampling at multiple time points was not possible. To get around this, we used sacrificial sampling by utilizing four to six replicates that were sequentially removed for measurement at various time points during the incubation period. For the time-series data, we assayed: pH, SCOD, TCOD, acetate, propionate, *n*-butyrate, $H_{2(g)}$, $CH_{4(g)}$, and $CO_{2(g)}$. However, because propionate and *n*-butyrate were only detected at very low concentrations, these data are not reported. Similarly, in most cases, $CO_{2(g)}$ comprised the majority of gas production with only trace levels of $H_{2(g)}$ and virtually no $CH_{4(g)}$. For this reason, we only report total biogas production, which was not converted to standard temperature and pressure (STP), for the preliminary experiments. Finally, to determine whether the fermentation product spectra were statistically different between test conditions, either a paired student *t*-test or repeated measures analysis of variance (rANOVA) in the car package in R Studio *v*3.4.4 was performed. The paired student *t*-test was used in experiments involving only two test conditions (*i.e.,* freezing *vs*. no freezing, mixing *vs*. no mixing, and inoculum *vs*. no inoculum), while the rANOVA was used in experiments involving three test conditions (*i.e.,* temperature and feeding strategy). A significance level of 0.05 was applied for both statistical approaches.

**Section S1.2: Effect of storage temperature**

Food waste was stored at three different storage temperature conditions (*i.e.,* 15°C, 25°C, and

37°C) to determine the effect of storage temperature on the fermentation process. Five replicate media bottles containing the same food waste were prepared for each of the three storage temperature conditions. The composition of the food waste used in this preliminary experiment is given in **Table S1**. The prepared bottles were then stored without mixing in a temperature-controlled room for an incubation period of 10 days. One bottle from each set was removed for sampling on Days: 1, 2, 3, 8, and 10.

**Table S1.** Characterization of the food waste used in the storage temperature experiment. BD = below detection

| **Variable** | **Value** | **Units** | ***n*** |
|---|---|---|---|
| Ammonia | 28 | mg $NH_3$-N $L^{-1}$ | 1 |
| Total solids | 119 ± 1 | g $L^{-1}$ | 3 |
| Volatile solids | 112 ± 1 | g $L^{-1}$ | 3 |
| pH | 5.55 | - | 1 |
| SCOD | 53 ± 2 | g COD $L^{-1}$ | 5 |
| TCOD | 109 ± 11 | g COD $L^{-1}$ | 3 |
| Acetate | 16 ± 220 | mg COD $L^{-1}$ | 2 |
| Propionate | BD | mg COD $L^{-1}$ | 2 |
| *n*-Butyrate | 65 ±18 | mg COD $L^{-1}$ | 2 |

Results of the experiment suggest that the fermentation process was affected by storage temperature, since the acetate concentration exhibited statistically significant differences between temperature conditions at one or more time points (**Fig. S2**). However, no statistical differences were found for any of the remaining variables.

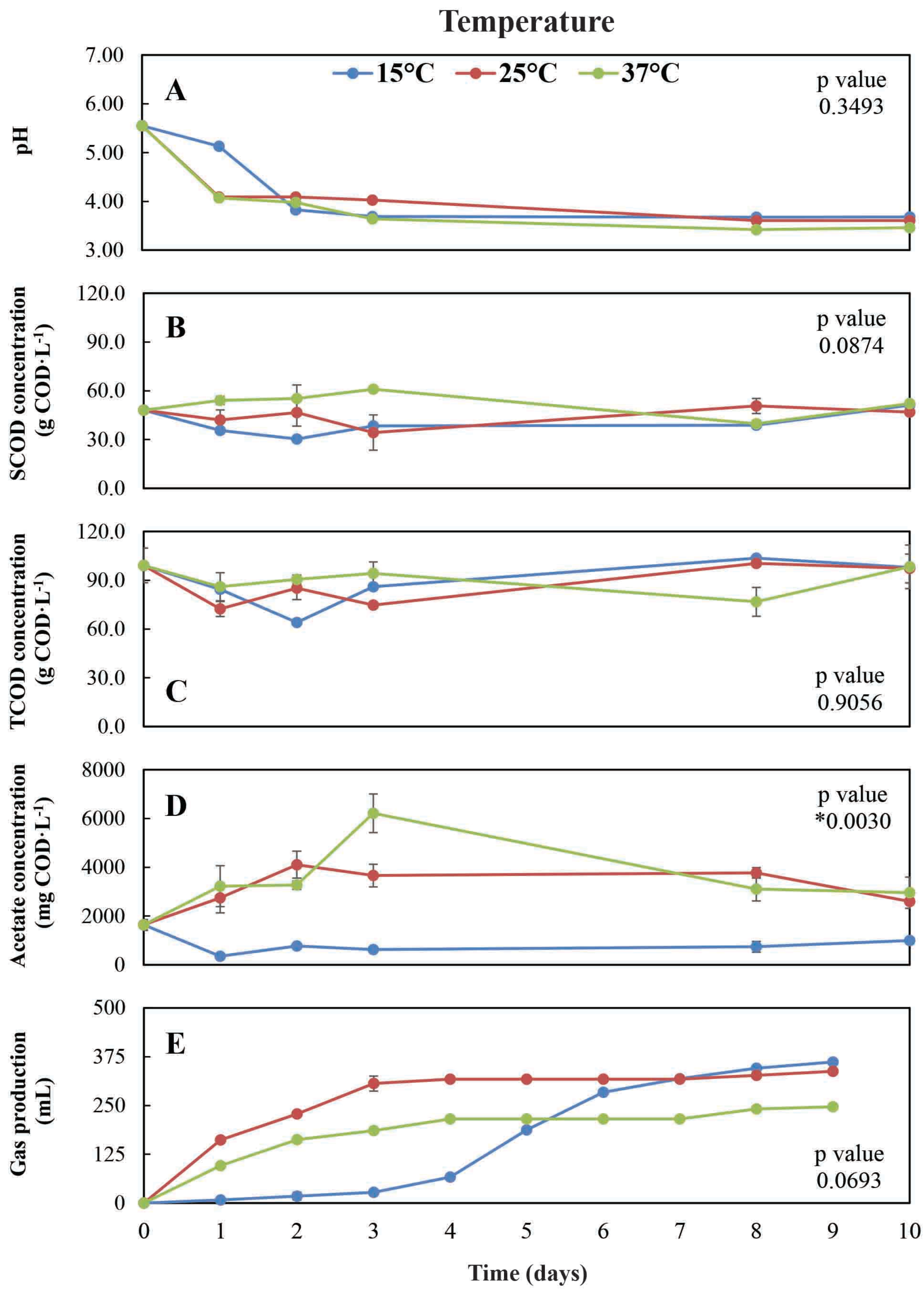


**Figure S2.** Time-series data collected during the storage temperature experiment involving three temperatures: 15°C (blue-line), 25°C (red-line), and 37°C (green-line). The resulting p values from the rANOVA analyses are indicated for each variable. The symbol '*' indicates a statistically significant p value.

**Section S1.3: Effect of feeding strategy**

In this preliminary experiment, food waste was added to the bottles using three different feeding strategies to determine their effects on the fermentation process. The first feeding strategy was single batch, where the bottles were fed once completely at the start of the incubation period. The second and third feeding strategies were semi-batch, where the bottles were periodically fed smaller amounts of food waste during the course of the incubation period. The volume added during each feeding cycle was equivalent to one-sixth of the final volume (*i.e.,* 30 mL). In the first semi-batch feeding strategy (*i.e.,* semi-batch: anoxic), the headspace was rigorously sparged with $N_2$ gas during feeding. However, in the second semi-batch strategy (*i.e.,* semi-batch: semi-oxic), the headspace was deliberately exposed to air for several min during and after feeding to determine the effect of oxygen exposure on the fermentation process. For each feeding strategy, a set of six replicate media bottles containing only food waste were prepared and incubated for 5 days at a temperature of 25°C without mixing. The composition of the food waste used in this preliminary experiment is given in **Table S2**. One bottle from each set was removed for sampling on Days: 1, 2, 3, 4, and 5.

**Table S2.** Characterization of the food waste used in the storage feeding strategy experiment. BD = below detection.

| Variable | Value | Units | *n* |
|---|---|---|---|
| Ammonia | 63 | mg $NH_3$-N $L^{-1}$ | 1 |
| Total solids | 203 ± 18 | g $L^{-1}$ | 3 |
| Volatile solids | 190 ± 14 | g $L^{-1}$ | 3 |
| pH | 5.89 | - | 1 |
| SCOD | 49 ± 7 | g COD $L^{-1}$ | 4 |
| TCOD | 124 ± 9 | g COD $L^{-1}$ | 5 |
| Acetate | 116 ± 52 | mg COD $L^{-1}$ | 2 |
| Propionate | BD | mg COD $L^{-1}$ | 2 |
| *n*-Butyrate | BD | mg COD $L^{-1}$ | 2 |

Results of the experiment suggest that the fermentation process was affected by the specific feeding strategy employed, since the SCOD concentration exhibited statistically significant differences between conditions at one or more time points (**Fig. S3**). However, no statistical differences were found for any of the remaining variables.

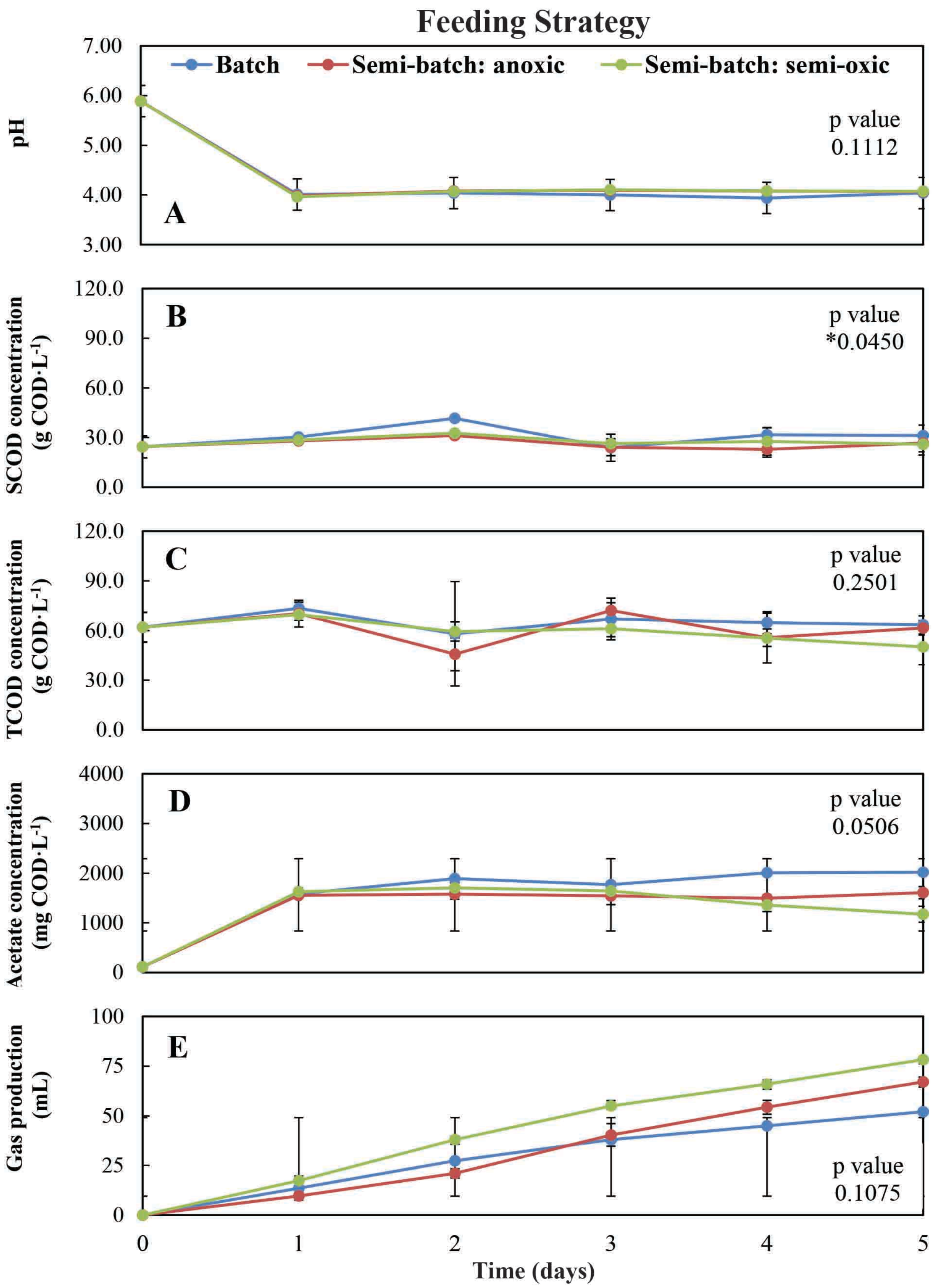


**Figure S3.** Time-series data collected during the storage feeding strategy experiment. Three feeding strategies were investigated: 1) batch-fed (blue line), 2) semi-batch fed with anoxic headspace (red line), and 3) semi-batch fed with semi-oxic headspace (green line). The resulting p values from the rANOVA analyses are indicated for each variable. The symbol '*' indicates a statistically significant p value.

### Section S1.4: Effect of inoculum presence

Here, food waste was stored in either the presence or absence of a fermentative inoculum to determine its effects on the fermentation process. The inoculum originated from an existing food waste storage tank that had been in operation for approximately 9 months. The inoculum was taken directly from the storage tank, which contained 6 days' worth of food waste. The composition of the food waste and inoculum used in this preliminary experiment is given in **Table S3**. Two sets of five replicate media bottles containing only food waste or food waste plus 10% (*v/v*) inoculum were incubated for 9 days at a temperature of 25°C without mixing. In addition, another set of five replicates containing only inoculum and water were used as controls. One bottle from each set was removed for sampling on Days: 1, 2, 3, 7, and 9.

**Table S3.** Characterization of the food waste and inoculum used in the inoculum experiment. BD = below detection.

| **Variable** | **Value** | | **Units** | ***n*** | |
|---|---|---|---|---|---|
| | **Inoculum** | **Food waste** | **-** | **Inoculum** | **Food waste** |
| Ammonia | 289 | 19 | mg $NH_3$-N $L^{-1}$ | 1 | 1 |
| Total solids | 105 ± 4 | 177 ± 14 | g $L^{-1}$ | 4 | 4 |
| Volatile solids | 97 ± 4 | 143 ± 13 | g $L^{-1}$ | 4 | 4 |
| pH | 3.96 | 3.73 | - | 1 | 1 |
| SCOD | 4 ± 0.15 | 49 ± 1 | g COD $L^{-1}$ | 3 | 3 |
| TCOD | 8 ± 1 | 100 ± 11 | g COD $L^{-1}$ | 4 | 5 |
| Acetate | 70 ± 24 | 272 ± 654 | mg COD $L^{-1}$ | 2 | 2 |
| Propionate | BD | 145 ± 35 | mg COD $L^{-1}$ | 2 | 2 |
| *n*-Butyrate | BD | 9 ± 0 | mg COD $L^{-1}$ | 2 | 2 |

Results of the experiment suggest that the fermentation process was affected by the presence of an inoculum. Both the pH and the acetate concentration exhibited statistically significant differences between conditions at one or more time points (**Fig. S4**). However, no statistical differences were found for any of the remaining variables.

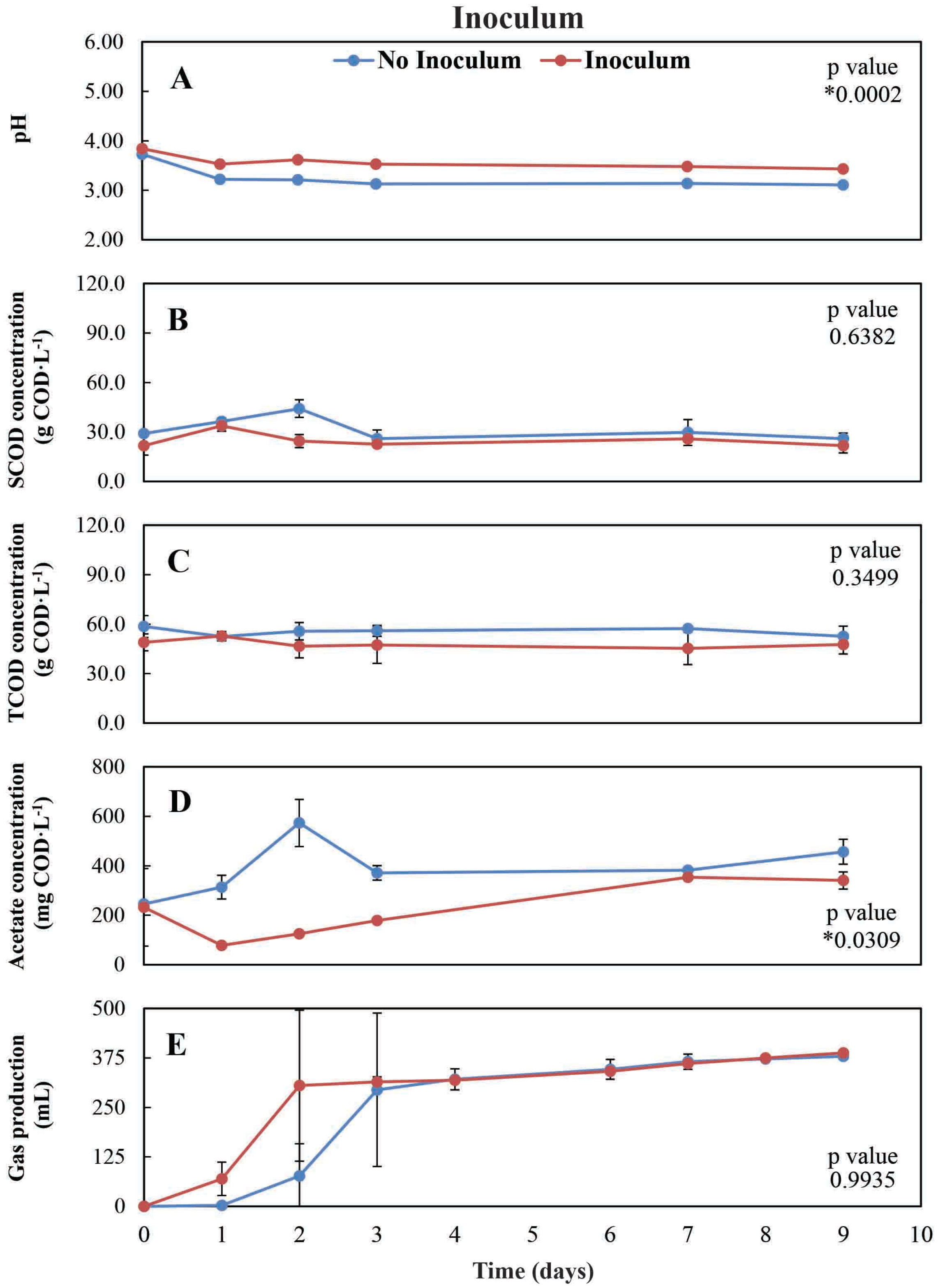


**Figure S4.** Time-series data collected during the storage inoculum experiment. Two test conditions were investigated: 1) food waste with no inoculum present (blue line), and 2) food waste with inoculum (red line). The resulting p values from the paired student *t*-test analyses are indicated for each variable. The symbol '*' indicates a statistically significant p value.

### Section S1.5: Effect of mixing

The objective of this preliminary experiment was to determine the effect of mixing on the fermentation process during the storage of food waste. Fresh food waste was collected, ground, and then stored at 4°C until use. Two sets of five replicate media bottles containing only food waste were prepared and incubated for 11 days at a temperature of 25°C. One set of replicates was not mixed, while the other set was manually mixed twice daily for the duration of the incubation period. The composition of the food waste used in this preliminary experiment is given in **Table S4**. One bottle from each set was removed for sampling on Days: 1, 2, 3, 7, and 11.

**Table S4.** Characterization of the food waste used in the storage mixing experiment. BD = below detection.

| **Variable** | **Value** | **Units** | ***n*** |
|---|---|---|---|
| Ammonia | 28 | mg $NH_3$-N $L^{-1}$ | 1 |
| Total solids | 33 | g $L^{-1}$ | 1 |
| Volatile solids | 28 | g $L^{-1}$ | 1 |
| pH | 5.05 | - | 1 |
| SCOD | 8 ± 0.5 | g COD $L^{-1}$ | 3 |
| TCOD | 11 ± 2 | g COD $L^{-1}$ | 2 |
| Acetate | 43±9 | mg COD $L^{-1}$ | 2 |
| Propionate | BD | mg COD $L^{-1}$ | 1 |
| *n*-Butyrate | BD | mg COD $L^{-1}$ | 1 |

Results of the experiment suggest that the fermentation process was not affected by mixing, since not one of the variables exhibited a significant difference between conditions at any time point (**Fig. S5**).

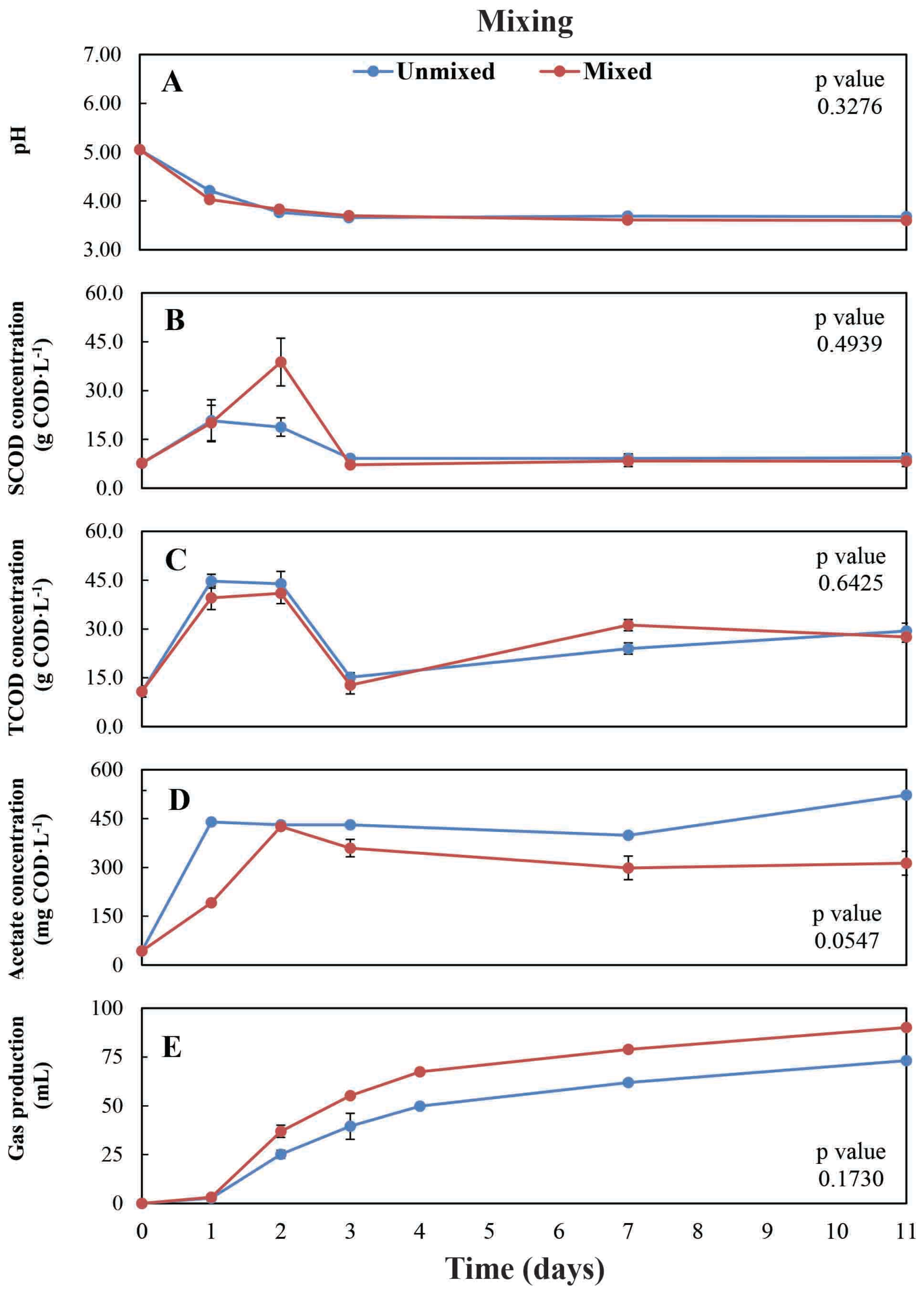


**Figure S5.** Time series data collected during the storage mixing experiment. The food waste was stored and either remained unmixed (blue line) or manually mixed (red line). The resulting p values from the paired student *t*-test analyses are indicated for each variable. The symbol '*' indicates a statistically significant p value.

**Section S1.6: Effect of freezing**

The objective of this preliminary experiment was to determine the effects of freezing food waste on the fermentation product spectrum during storage. A single batch of fresh food waste was collected, ground, and then split into two smaller batches. One of the small batches was kept in a refrigerator at 4°C, while the other was kept in a freezer at -20°C for approximately 24 h. On the day the bottles were prepared, each small batch of food waste was removed from cold storage and left to equilibrate to room temperature. Then, two sets of five replicate media bottles containing either the fresh or previously frozen food waste were prepared and incubated for 9 days at a temperature of 25°C without mixing. The composition of the food waste used in this preliminary experiment is given in **Table S5**. One bottle from each set was removed for sampling on Days: 1, 2, 3, 7, and 9.

**Table S5.** Characterization of the food waste used in the storage freezing experiment. BD = below detection.

| **Variable** | **Value** | **Units** | ***n*** |
|---|---|---|---|
| Ammonia | 8 | mg $NH_3$-N $L^{-1}$ | 1 |
| Total solids | 241 ± 2 | g $L^{-1}$ | 4 |
| Volatile solids | 194 ± 4 | g $L^{-1}$ | 4 |
| pH | 6.27 | - | 1 |
| SCOD | 33 ± 5 | g COD $L^{-1}$ | 5 |
| TCOD | 219 ± 14 | g COD $L^{-1}$ | 5 |
| Acetate | 1940 ± 90 | mg COD $L^{-1}$ | 2 |
| Propionate | BD | mg COD $L^{-1}$ | 2 |
| *n*-Butyrate | BD | mg COD $L^{-1}$ | 2 |

Results of the experiment suggest that the fermentation process was not affected by freezing, since not one of the variables exhibited a significant difference between conditions at any time point (**Fig. S6**).

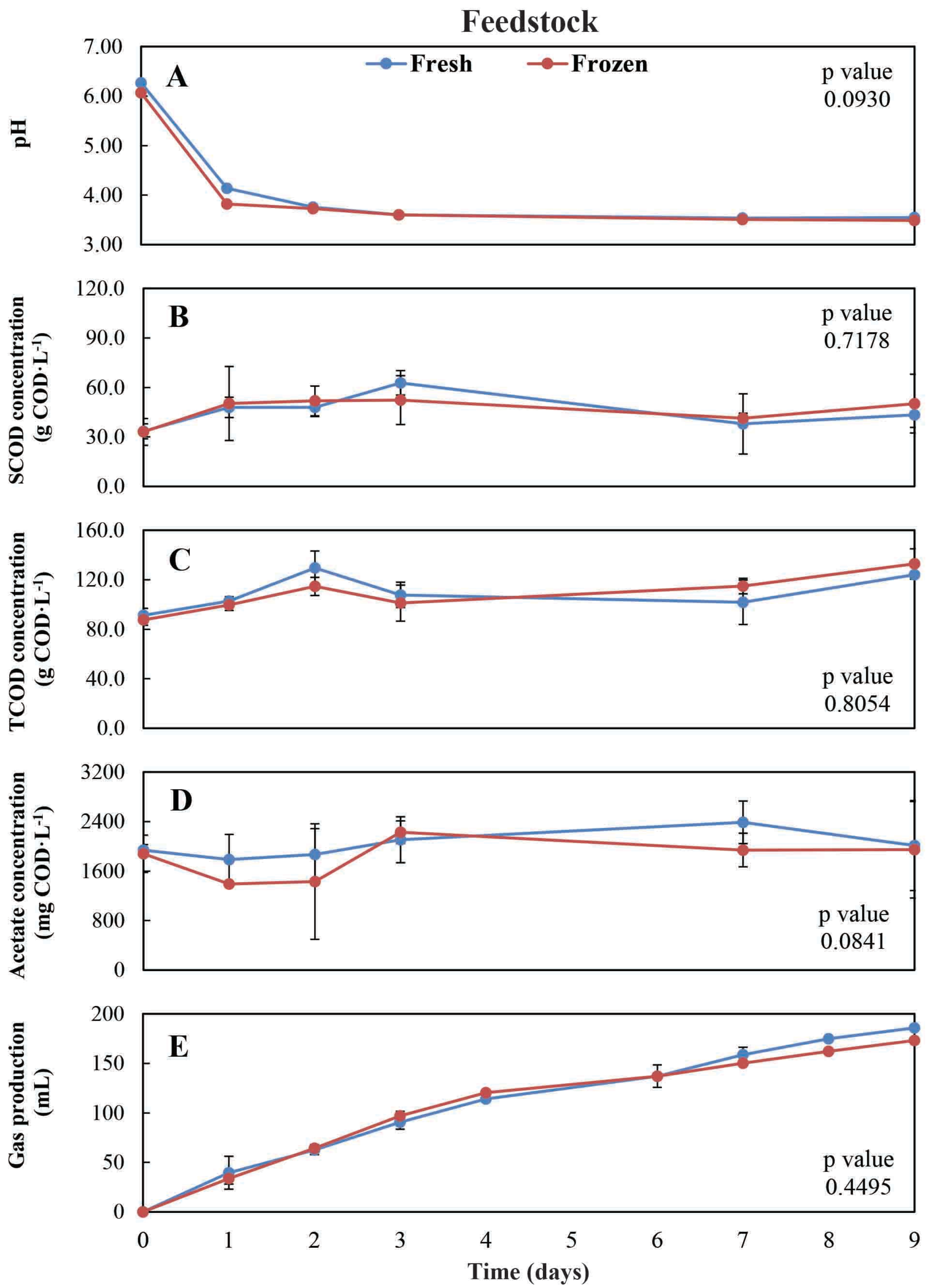


**Figure S6.** Time series data collected during the storage freezing experiment. The food waste was used as either fresh (blue line) or frozen feedstock (red line) for the bottle experiment. The resulting p values from the paired student *t*-test analyses are indicated for each variable. The symbol '*' indicates a statistically significant p value.

**III) Fermentation product spectrum comparison for the two protein-rich feedstocks for the storage experiment**

We made a mistake during collection of food waste for the storage experiment and did not obtain enough protein-rich food waste. Therefore, a second collection and preparation event was performed. The analyses showed that the pH was similar between the two different protein-rich food-waste batches (6.4 and 6.6 in **Table 3**); however, the TCOD concentration for the protein-rich I food waste was higher than for the protein-rich II food waste (67 and 38 g $L^{-1}$, respectively in **Table 3**), which did not result in a large difference in the ratio of SCOD $TCOD^{-1}$ (21 and 26, respectively, in **Table 3**). We compared the fermentation product spectra for the two different protein-rich food-waste batches within the individual experimental combinations (**Table 1**). We found no significant differences ($p>0.53$) in the fermentation product spectra (pH, ammonia concentration, $CO_2$ volume, and $H_2$ volume) between three large-media bottles during the experimental period for the two protein-rich food-waste batches (**Fig. S7A-D**). We are, therefore, confident that the two different protein-rich food-waste batches behaved similarly enough in the storage experiment to permit the statistical analysis.

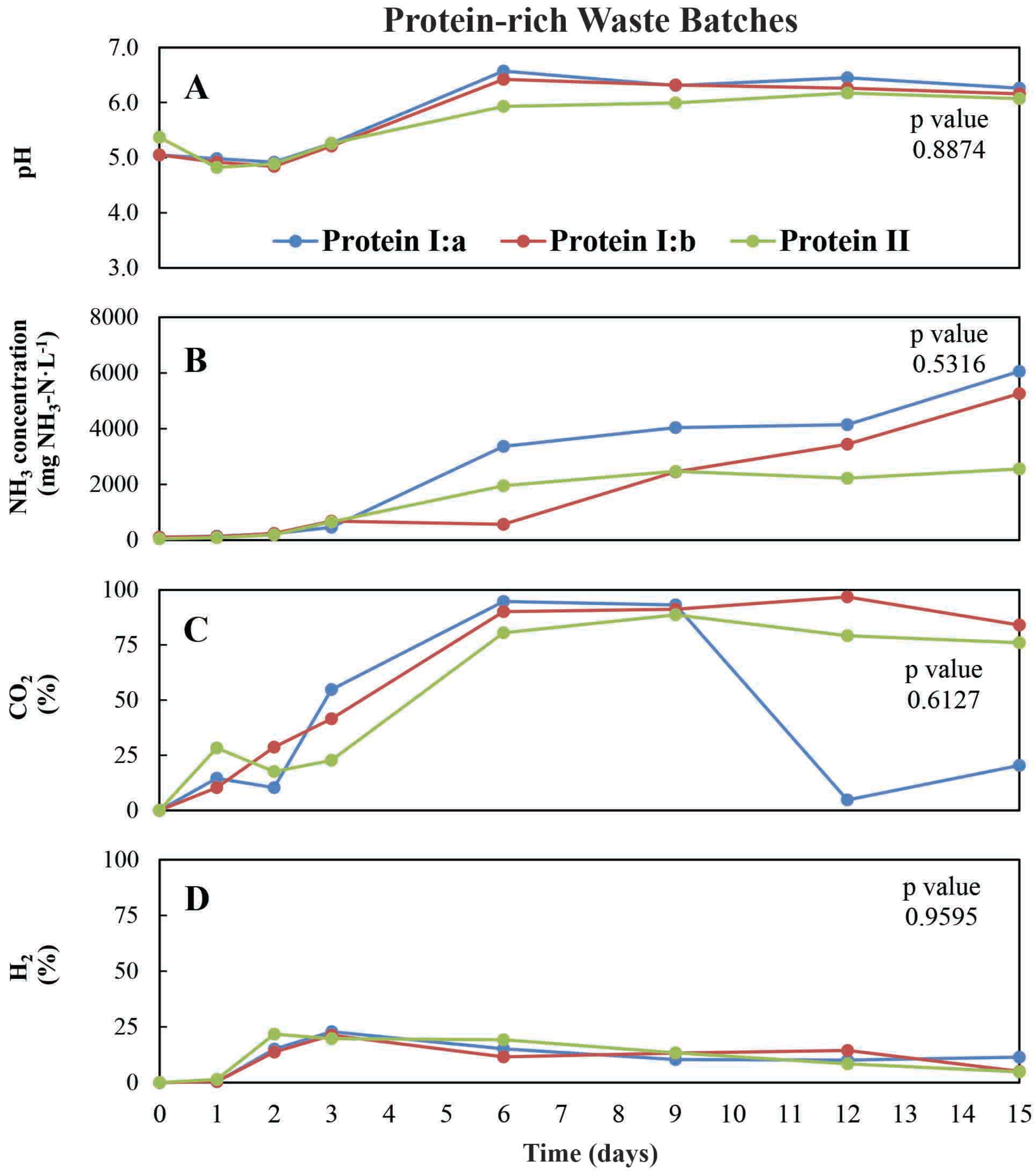


**Figure S7.** Time series data collected during the large-media-bottle experiment to investigate whether the protein-rich I and the protein-rich II food-waste batches acted significantly different. This data includes a duplicate protein-rich I food waste (blue and red line) and a single protein-rich II food waste (green line). The resulting p values from the rANOVA analyses are indicated for each variable. We did not observe a statistically significant difference.

### IV) MANOVA of data from the storage experiment

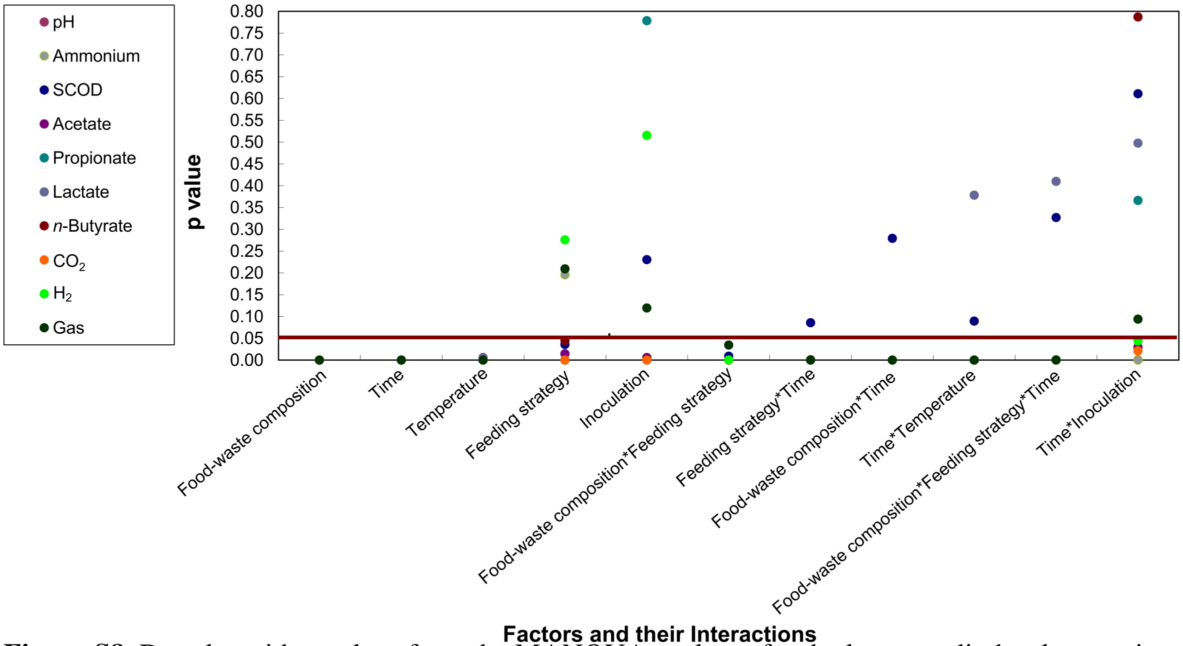


**Figure S8.** Dot plot with p values from the MANOVA analyses for the large-media-bottle experiment. We performed MANOVA analysis with the entire dataset including each dependent variables (pH, ammonia concentration, SCOD concentration, acetate concentration, propionate concentration, lactate concentration, *n*-butyrate concentration, $CO_2$ production, $H_2$ production, and gas production). The p values indicate whether the different factors *plus* time in the experiment (storage temperature, food-waste composition, feeding strategy, inoculation, and time) or their two-way and three-way interactions significantly ($p<0.05$) influenced the dependent variable. The factors and their interactions are ordered from low to high p value. Values above 0.05 (red line) indicate that the dependent variable was not significantly influenced by the factor. We only included significant interactions on this graph. Note that for values below 0.05, some dots overlap.

### V) Principle component analysis (PCA) of data from the storage experiment

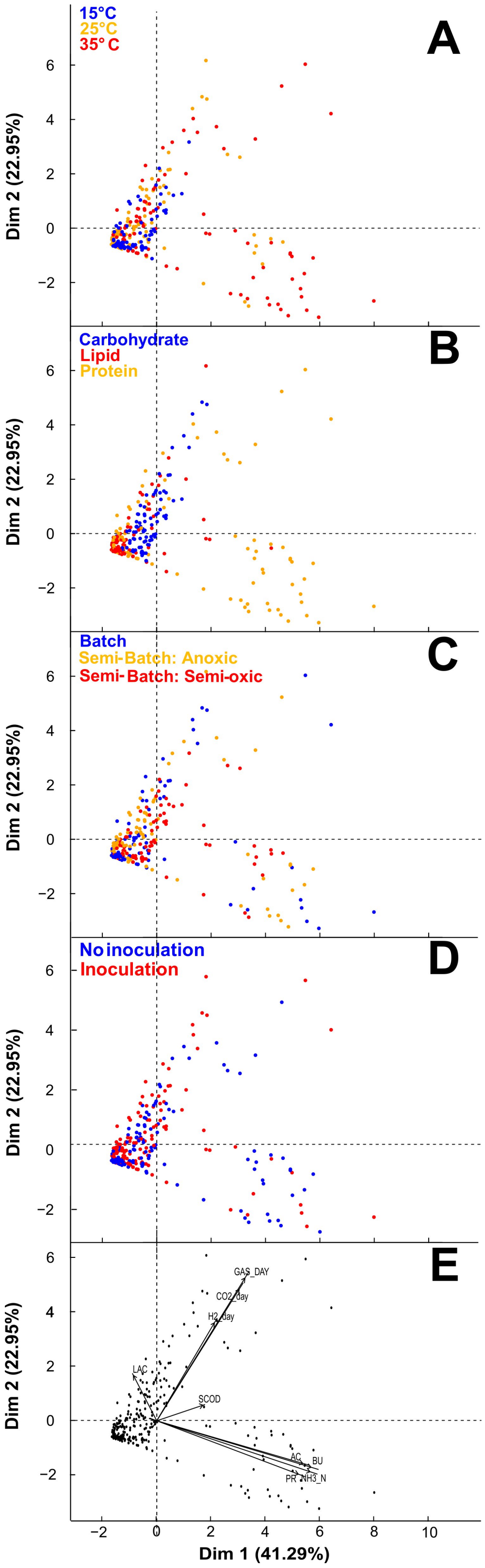

**Figure S9.** Principle Component Analysis plots of storage bottles (288 sampling points from the large-media bottles for all days and sampling conditions) throughout the 15-day operating period. Two principle components are shown. Dim 1 explained 41.29% of the variation and Dim 2 explained 22.95% of the variation. **A.** The colors of each circle represent the temperature of the experiment (blue = 15°C; orange = 25°C; and red = 35°C). **B.** The colors of each circle represent the food-waste composition of the bottle (blue = carbohydrate-rich; red = lipid-rich; and orange = protein-rich). **C.** The colors of each circle represent the feeding strategy (blue = batch = no periodic feed; orange = semi—batch anoxic = period feed, but no oxygen exposure; and red = semi-batch semi-oxic = periodic feed, with oxygen exposure). **D.** The colors of each circle represent the inoculation (blue = no inoculation; and red = inoculation). **E.** Biplot of the 288 sampling points with each vector (line with arrow) representing a dependent variable. The angle between two vectors represents the correlation between those particular variables.

### VI) Interaction plots for the factor feeding strategy *plus* time from the storage experiment

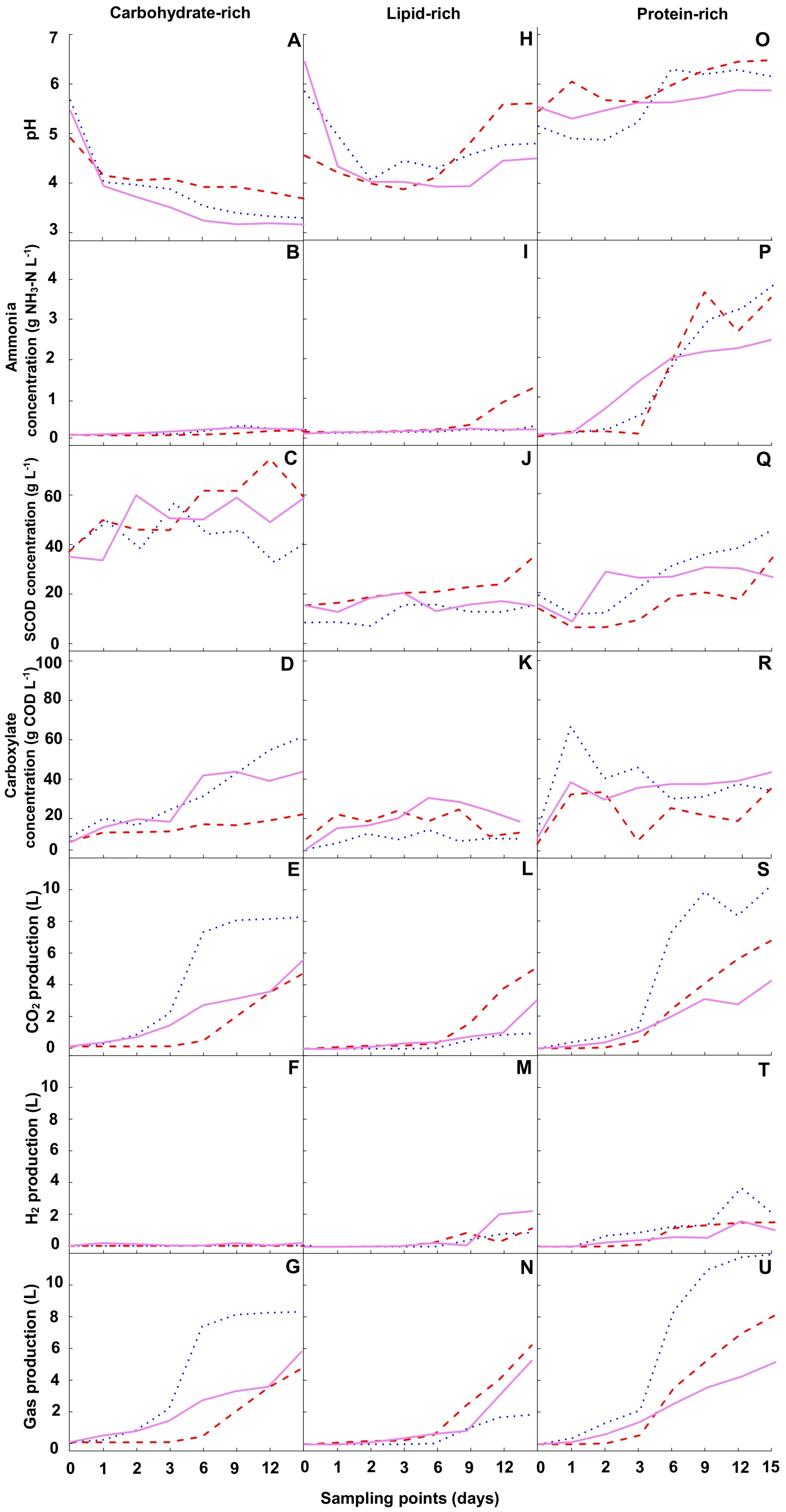


**Figure S10.** Twenty-one interaction plots for each dependent variable and for each of three food-waste compositions throughout the operating period. Each of these plots is made with one of three different linear regression models (one for each food-waste composition) with the interacting factors feeding strategy *plus* time. The resulting regression line with the response variables represents the predicted daily dependent variable throughout the operating period of the large-media-bottle-experiment for one feeding strategy with a total of three lines for each feeding strategy (blue dotted line = Batch; violet solid line = Semi-Batch: Anoxic; and red dashed line = Semi-Batch: Semi-oxic). Here, we show seven different dependent variables (pH, ammonia concentration, SCOD concentration, total carboxylate concentration, $CO_2$ production, $H_2$ production, and total gas production) throughout the operating period for three different types of food waste (carbohydrate-rich, lipid-rich, and protein-rich), resulting in 21 panels (**A-U**). Curves that are not parallel within one panel indicate interaction between feeding strategy and time. Overall, changes in feeding strategy do not have a linear effect on the dependent variable. Note that the x-axis (time) is not to scale on purpose to identify that this is data based on a model.

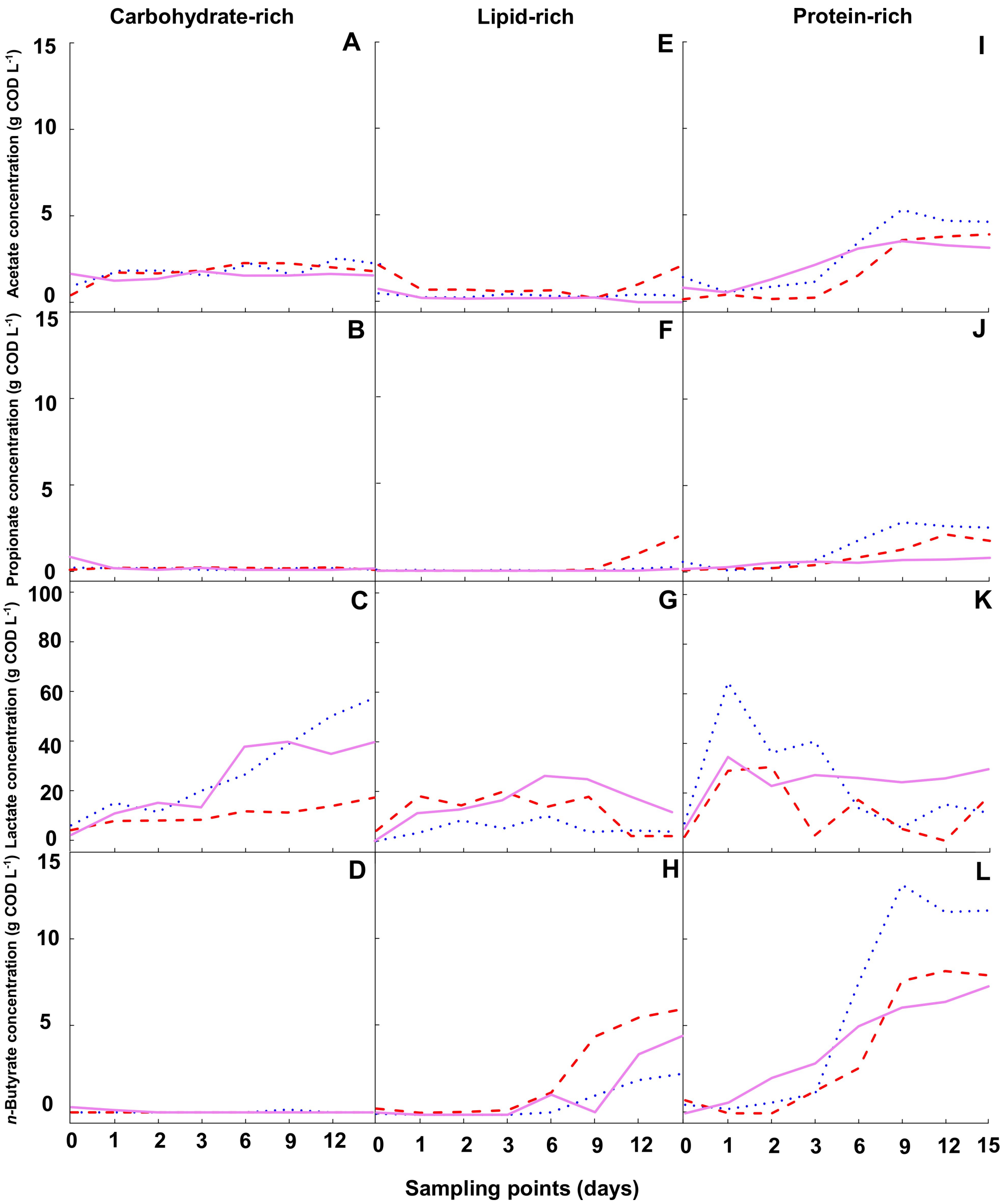


**Figure S11.** Twelve interaction plots for each of the SCC concentrations (dependent variables) throughout the operating period. The same linear regression models with the interacting factors feeding strategy *plus* time were used as in **Fig. S10**. Three lines for each temperature (blue dotted line = Batch; violet solid line = Semi-Batch: Anoxic; and red dashed line = Semi-Batch: Semi-oxic) in each panel show the predicted SCC concentrations for acetate, *n*-propionate, lactate, and *n*-butyrate throughout the operating period for each temperature. With three different types of food waste (carbohydrate-rich, lipid-rich, and protein-rich), this resulted in 12 panels (**A-L**). Curves that are not parallel within one panel indicate interaction between feeding strategy and time. Overall, changes in feeding strategy do not have a linear effect on the SCC concentrations. Note that the x-axis (time) is not to scale on purpose to identify that this is data based on a model.

### VII) Interaction plots for the factor inoculation *plus* time from the storage experiment

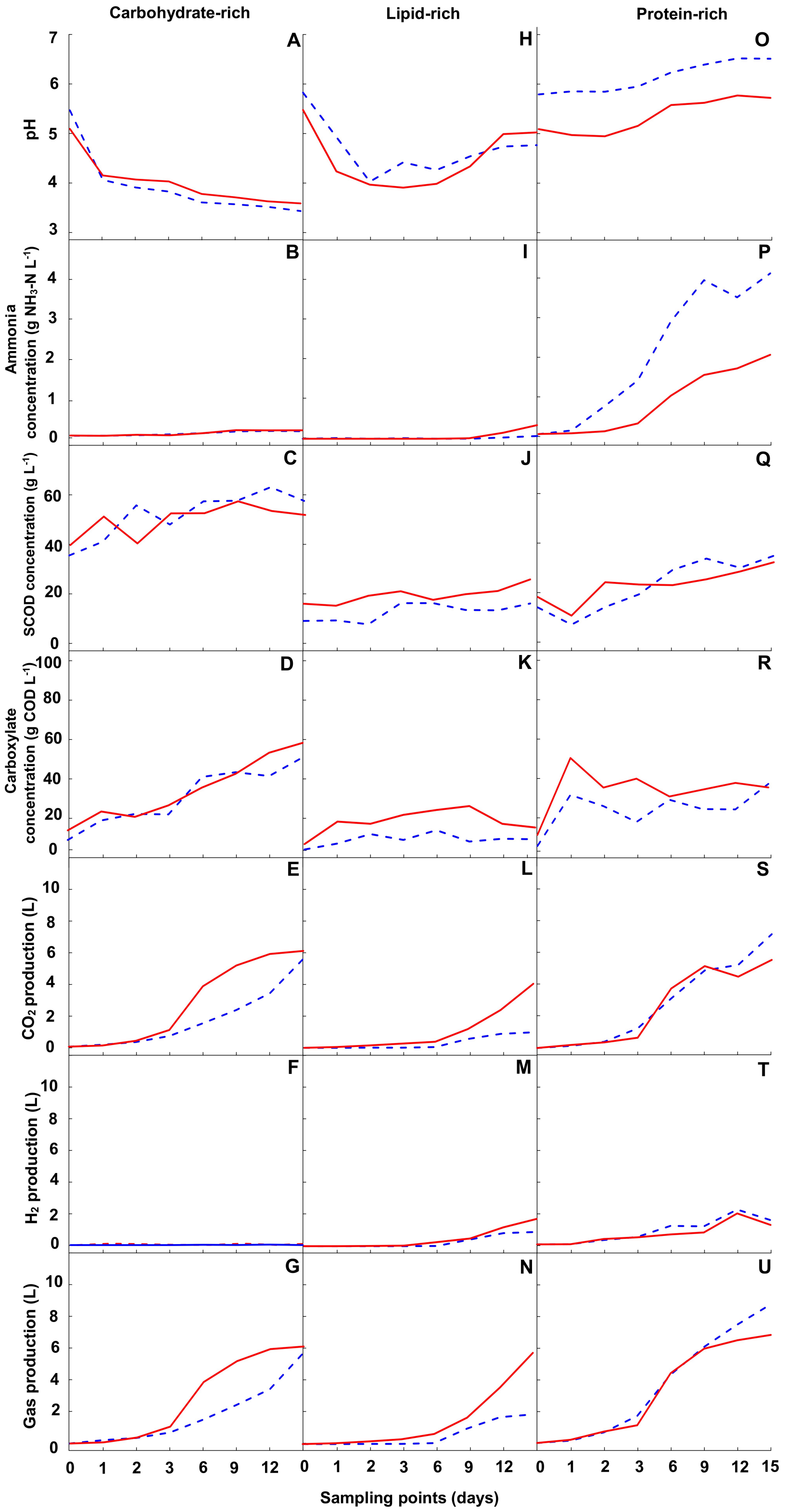


**Figure S12.** Twenty-one interaction plots for each dependent variable and for each of three food-waste compositions throughout the operating period. Each of these plots is made with one of three different linear regression models (one for each food-waste composition) with the interacting factors inoculation *plus* time. The resulting regression line with the response variables represents the predicted daily dependent variable throughout the operating period of the large-media-bottle experiment for one feeding strategy with a total of two lines for each feeding strategy (red solid line = Yes; blue dashed line = No). Here, we show seven different dependent variables (pH, ammonia concentration, SCOD concentration, total carboxylate concentration, $CO_2$ production, $H_2$ production, and total gas production) throughout the operating period for three different types of food waste (carbohydrate-rich, lipid-rich, and protein-rich), resulting in 21 panels (**A-U**). Curves that are not parallel within one panel indicate interaction between inoculation and time. Overall, changes in inoculation do not have a linear effect on the dependent variable. Note that the x-axis (time) is not to scale on purpose to identify that this is data based on a model.

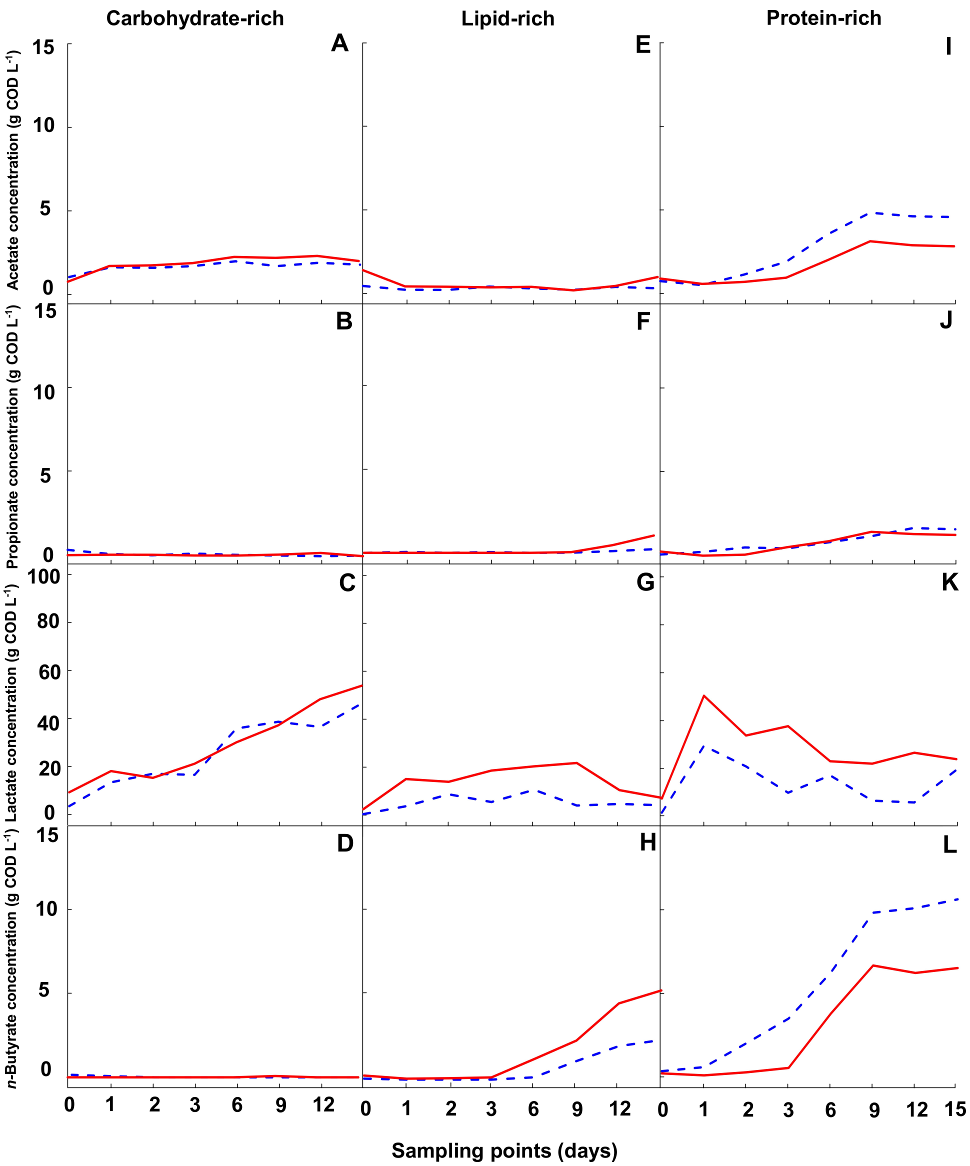


**Figure S13.** Twelve interaction plots for each of the SCC concentrations (dependent variables) throughout the operating period. The same linear regression models with the interacting factors feeding strategy *plus* time were used as in **Fig. S12**. Three lines for each temperature (red solid line = Yes; blue dashed line = No) in each panel show the predicted SCC concentrations for acetate, *n*-propionate, lactate, and *n*-butyrate throughout the operating period for each temperature. With three different types of food waste (carbohydrate-rich, lipid-rich, and protein-rich), this resulted in 12 panels (**A-L**). Curves that are not parallel within one panel indicate interaction between feeding strategy and time. Overall, changes in feeding strategy do not have a linear effect on the SCC concentrations. Note that the x-axis (time) is not to scale on purpose to identify that this is data based on a model.

**<u>VIII) Linear regression equations for lactate and $H_2$ for the factors *plus* time from the storage experiment</u>**

The following linear regression equation depicts the relationship between lactate concentration and the factors time, temperature, food-waste composition, feeding strategy, and inoculation (from our model with the units):

**LACTATE = 9.3 – 2.06*TIME + 3.27*TEMPERATURE – 4.17*FOOD-WASTE COMPOSITION + 3.91*FEEDING STRATEGY + 9.34*INOCULATION (S1)**

where the units for: lactate = g COD $L^{-1}$; and time = days. For qualitative factor levels: temperature (15°C = 1; 25°C = 3; and 35°C = 2); food-waste composition (carbohydrate-rich = 1; lipid-rich = 2; and protein-rich = 3); feeding strategy (batch = 1; semi-batch, semi-oxic = 2; and semi-batch: anoxic = 3); and inoculation (no = 1; and yes = 2).

In our study, a linear regression equation with only temperature and food-waste composition depicted the relationship between $H_2$ production and the factors (from our model with the units):

**$H_2$ = -81.62 + 27.91*TEMPATURE + 40.47*FOOD-WASTE COMPOSITION (S2)**

where the units for: $H_2$ = mL. For qualitative factor levels: temperature (15°C = 1; 25°C = 3; and 35°C = 2); and food-waste composition (carbohydrate-rich = 1; lipid-rich = 2; and protein-rich = 3).

**IX) BMP experiment to evaluate methane production from stored food waste from the storage experiment.**

In **Section S9.1**, we provide a description of the general methodology that was applied for BMP experiment, while in **Sections S9.2** we explain the results in detail.

**Section S9.1: General approach for the BMP experiments**

We combined the BMP inoculum with a sample from the storage experiment at a VS mixture ratio of 1:1 (*w/w*), respectively. For the BMP inoculum, we used fresh anaerobic digestion sludge collected from the Ithaca Area Wastewater Treatment Facility (Ithaca, New York), which we first acclimated with a food-waste sample with a 1:1:1 ratio (*w/w*) of carbohydrate, lipid, and protein at a temperature of 25°C for two weeks. The VS concentration of the BMP inoculum was 5.11 ± 0.34 g VS $L^{-1}$. We then added a fixed volume of a nutrient and buffer medium (pH ranged between 7.1 and 7.3), and a balance of deoxygenated water to achieve a final volume of 100 mL. The final sample concentration was 1 g VS $L^{-1}$. The BMP mixture was then added to 250-mL glass media bottles, sealed, and then sparged with $N_2$ gas. Next, the bottles were placed in incubation chambers set to 37±1°C where they were kept for at least 30 days or until the cumulative biogas production had leveled off.

During incubation, we used the gas-displacement method, using a 30-mL glass syringe (Chemglass Life Sciences Co.) to measure the gas production. Immediately after measuring biogas production, we measured biogas composition (*i.e.,* $N_2$, $CH_4$, and $CO_2$). To account for endogenous biogas production from the inoculum, we prepared blank BMP mixtures containing only BMP inoculum and nutrient media. The biogas produced from these blank BMPs was then subtracted from the total biogas measured in the BMPs containing the sample substrate. Finally, biogas production volumes were normalized to standard conditions, similar to the storage experiment.

**Section S9.2: BMP experiment results**

Because we had not observed a significant decrease ($p$ value < 0.05) in TCOD concentrations during food storage (**Table S6**), we normalized the methane production based on the TCOD concentration for the pre-storage (fresh) food waste or the post-storage food waste. Others have used VS concentration for normalization (Labatut et al. 2011), however, during food-waste storage the VS concentrations had decreased while the soluble fermentation products increased considerably during hydrolysis and acidification (**Table S6**), resulting in a biased normalization. In general, TCOD concentration measurements from a slurry with solids are recognized to have relatively large error due to: 1) a pipetting error due to the very large dilutions that are necessary; and 2) a sampling error due to the non-homogeneous solid distribution after a sizeable dilution. Therefore, only large differences in methane selectivities became significantly different for our BMP data (**Fig. S12**).

We observed a significant increase in methane selectivity for BMP with a p value <0.01 for carbohydrate-rich food waste that was stored at 25°C compared to without storage (**Fig. S12**). This considerable increase was observed for the same large-media bottle with the maximum lactate concentration of 83 g COD $L^{-1}$ that we observed in this study (Comb. 5 in **Table S6**). At 35°C, we also observed a significant increase for carbohydrate-rich food waste, albeit at a higher p value <0.05 (**Fig. S12**). Finally, a significant methane selectivity increase was observed at 35°C for protein-rich food waste after storage compared to before storage for the BMP (p value <0.01 in **Fig. S12**). In all three of these large-media bottles, the lactate concentration was also high on Day 15 (Comb. 5, 9, and 12 in **Table S6**). Thus, the conversion of food waste into lactate during storage would be a positive for further methane generation in an anaerobic digestion system. It is unclear why other conditions, such as the protein-rich food waste at 15°C (Comb. 4 in **Table S6**), with similarly high lactate concentrations did not result in a significant increase in methane selectivity (**Fig. S12**). This condition was characterized by a very low ammonia concentration, and therefore a possible ammonia inhibition to methanogenesis could not explain the absence of the significant increase in methane selectivity (Rajagopal et al. 2013). For all combinations, we only observed a significant decrease in the methane selectivity for the BMP with stored *vs*. fresh carbohydrate-rich food waste at 15°C (p=0.048 in **Fig. S12**). However, this decrease cannot be explained by a toxic effect on methanogens due to accumulating breakdown products such as ammonia or LCFAs. In general, we found that food storage does not have a negative effect on methane formation in an anaerobic digestion system, however, for very specific conditions a significant lower methane selectivity may occur.

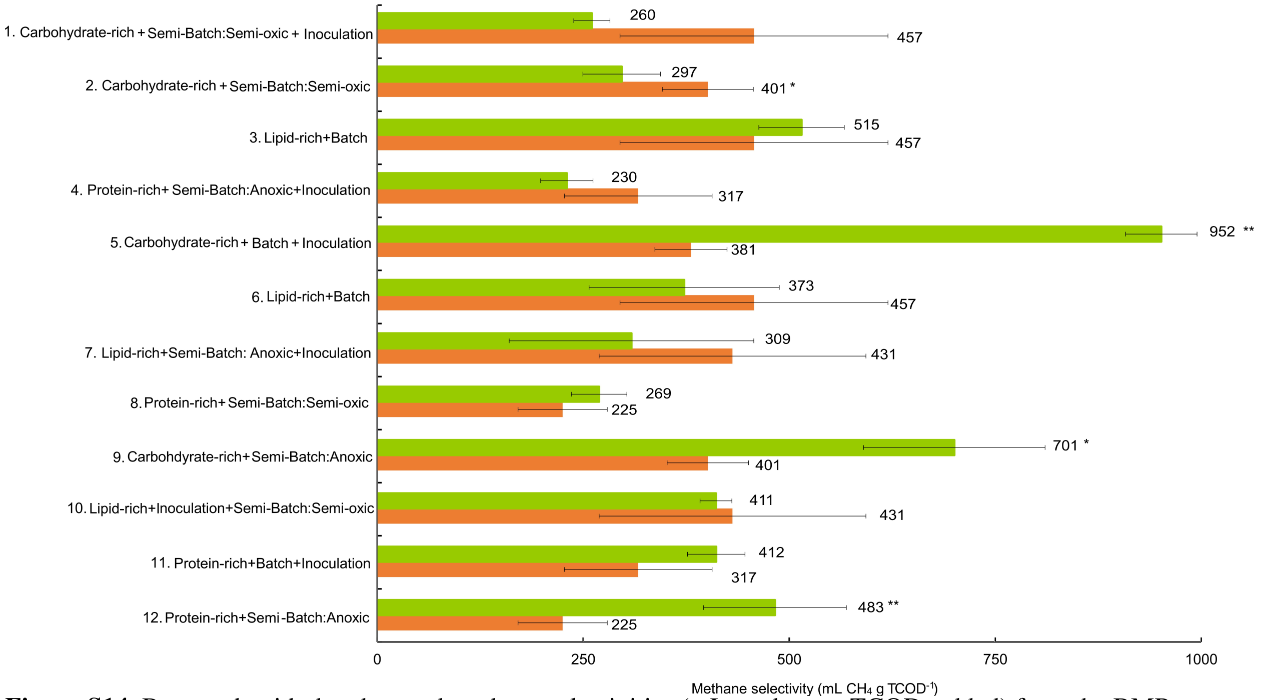


**Figure S14.** Bar graph with the observed methane selectivities (mL methane g $TCOD^{-1}$ added) from the BMP test with stored (green bar: post-storage) food waste and fresh food waste (red bar: pre-storage). The substrates were taken from the 12 different storage combinations on Day 0 (pre-storage) and Day 15 (post-storage). Batch or semi-batch refers to the feeding strategy of feedstock; anoxic or oxic refers to with or without removal of $O_2$ during the semi-batch feeding strategy; with or without inoculation refers to whether inoculum was added or not. The 12 combinations are separated by the temperature of the storage conditions (15°C – top four, 25°C – middle four, and 35°C – bottom four combinations). Significance codes: * p value<0.05, **p value<0.01, ***, p value<0.001.

### X) Measured data for each of the 12 experimental combinations from the storage experiment

**Table S6.** Measured data at the start (Day 0) and/or at the end (Day 15) of the storage experiment. When available or meaningful, we report the average ± standard deviation for triplicate experimental combinations. BD = below detection level; NA = data not available.

| Comb. | Day | pH | Ammonium conc. | VS conc. | TS conc. | VS $TS^{-1}$ | SCOD conc. | TCOD conc. | SCOD $TCOD^{-1}$ | Total carboxy-late conc. | Acetate conc. | Propionate conc. | Lactate conc. | Lactate Selectivity | *n*-Butyrate conc. | $CO_2$ prod. | $H_2$ prod. | Gas prod. |
|---|---|---|---|---|---|---|---|---|---|---|---|---|---|---|---|---|---|---|
| | | - | *(g $NH_3$ -N $L^{-1}$)* | *g $L^{-1}$* | *g $L^{-1}$* | *%* | *g COD $L^{-1}$* | *g COD $L^{-1}$* | *%* | *g COD $L^{-1}$* | *g COD $L^{-1}$* | *g COD $L^{-1}$* | *g COD $L^{-1}$* | *%* | *g COD $L^{-1}$* | *L* | *L* | *L* |
| 1 | 0 | 4.4±0.1 | 0.08±0.02 | 95.3 | 100 | 95 | 39 | 92 | 43 | 9.6 | 0.49 | BD | 9.1 | NA | BD | NA | NA | NA |
| 1 | 15 | 3.7±0.1 | 0.19±0.01 | 106±8.73 | 112±10.8 | 95±1.4 | 60±3.9 | 94±12 | 65±4.3 | 28±1.7 | 1.9±1.4 | BD | 26±1.4 | 29±1.5 | BD | 4.50±0.44 | BD | 4.50±0.44 |
| 2 | 0 | 5.4±0.3 | 0.07±0.03 | 95.6 | 100 | 96 | 35 | 92 | 38 | 4.0 | 0.43 | BD | 3.5 | NA | BD | NA | NA | NA |
| 2 | 15 | 3.7±0.1 | 0.16±0.01 | 114±2.31 | 119±3.40 | 95±0.8 | 59±30 | 99±19 | 64±33 | 26±3.4 | 1.8±1.1 | BD | 24±2.3 | 26±2.5 | BD | 4.87±2.77 | BD | 4.87±2.77 |
| 3 | 0 | 5.9±0.1 | 0.03±0.01 | 92.2 | 100 | 92 | 9±0 | 57 | 15 | 0.8 | 0.25 | BD | 0.57 | NA | BD | NA | NA | NA |
| 3 | 15 | 4.3±0.1 | 0.09±0.01 | 51.3±6.69 | 57.4±7.24 | 89±0.8 | 19±1.2 | 44±2.7 | 33±2.1 | 9.4±1.1 | 0.52±0.37 | BD | 8.8±1.4 | 15±2.4 | BD | 0.03±0.03 | BD | 0.03±0.03 |
| 4 | 0 | 5.0±0.1 | 0.05±0.01 | 38.6 | 100 | 39 | 17 | 44 | 39 | 8.0 | 0.38 | BD | 7.6 | NA | BD | NA | NA | NA |
| 4 | 15 | 5.3±0.0 | 0.23±0.03 | 43.0±8.71 | 55.5±18.0 | 79±9.9 | 18±0.8 | 50±3.3 | 42±1.7 | 37±41 | 1.1±0.09 | BD | 35±41 | 79±92 | 1.4±0.25 | 0.89±0.89 | 0.51±0.51 | 1.40±1.40 |
| 5 | 0 | 5.7±0.0 | 0.07±0.00 | 95.3 | 100 | 95 | 39 | 92 | 43 | 10 | 1.1 | 0.12 | 9.1 | NA | BD | NA | NA | NA |
| 5 | 15 | 3.3±0.0 | 0.20±0.05 | 75.3 | 70.4±9.09 | 93 | 41±23 | 91±46 | 45±25 | 85±12 | 2.2±0.16 | BD | 83±12 | 90±13 | BD | 8.28±1.89 | BD | 8.28±1.89 |
| 6 | 0 | 5.8±0.0 | 0.20±0.00 | 92.2 | 100 | 92 | 9 | 57 | 15 | 1.5 | 0.76 | BD | 0.57 | NA | 0.12 | NA | NA | NA |
| 6 | 15 | 5.3±0.2 | 0.37±0.30 | 42.4 | 41.9±10.5 | 86 | 13±2.2 | 51±14 | 23±3.9 | 8.2±3.3 | 0.22±0.38 | 0.44±0.40 | 2.9±2.6 | 5.0±4.5 | 4.7±0.79 | 1.93±0.80 | 1.77±0.07 | 3.71±0.79 |
| 7 | 0 | 6.4±0.0 | 0.02±0.00 | 92.5 | 100 | 93 | 16 | 61 | 26 | 1.5 | 0.76 | BD | 0.57 | NA | 0.13 | NA | NA | NA |
| 7 | 15 | 4.5±0.3 | 0.13±0.05 | 33.9 | 42.8±26.6 | 46 | 16±8.0 | 79±45 | 26±13 | 22±9.1 | BD | 0.10±0.18 | 17±8.3 | 28±14 | 4.5±1.3 | 3.01±0.75 | 2.24±0.39 | 5.25±1.03 |
| 8 | 0 | 5.5±0.0 | 0.02±0.00 | 71.5 | 100 | 72 | 14 | 67 | 21 | 2.5 | 0.20 | BD | 1.5 | NA | 0.75 | NA | NA | NA |
| 8 | 15 | 6.5±0.2 | 3.54±0.53 | 20.8 | 71.3±71.1 | 84 | 35±12 | 76±14 | 52±18 | 31±28 | 3.9±0.46 | 1.7±0.23 | 17±29 | 25±43 | 7.9±0.72 | 6.81±3.12 | 1.52±0.60 | 8.32±3.72 |
| 9 | 0 | 5.5±0.0 | 0.07±0.00 | 95.6 | 100 | 96 | 35 | 92 | 38 | 6.3 | 1.7 | 0.75 | 3.5 | NA | 0.30 | NA | NA | NA |
| 9 | 15 | 3.2±0.1 | 0.21±0.01 | 83.1±5.15 | 82.7±7.26 | 97±0.9 | 59±6.6 | 84±12 | 64±7.2 | 59±2.2 | 1.6±0.18 | 0.08±0.14 | 57±2.2 | 62±2.4 | BD | 5.56±0.09 | 0.20±0.34 | 5.75±0.25 |
| 10 | 0 | 4.5±0.0 | 0.07±0.00 | 92.5 | 100 | 93 | 16 | 61 | 26 | 8.9 | 2.1 | BD | 6.4 | NA | 0.35 | NA | NA | NA |
| 10 | 15 | 5.6±0.2 | 1.22±0.32 | 15.3±2.56 | 25.7±3.49 | 59±1.9 | 36±7.2 | 46±7.3 | 58±12 | 13±5.5 | 2.1±0.62 | 2.0±1.5 | 3.3±2.6 | 5.5±4.3 | 6.0±1.6 | 5.07±1.37 | 1.16±0.78 | 6.22±2.11 |
| 11 | 0 | 5.2±0.2 | 0.09±0.03 | 62.1±20.5 | 100 | 62 | 19 | 61 | 32 | 9.2 | 1.5 | 0.49 | 6.7 | NA | 0.49 | NA | NA | NA |
| 11 | 15 | 6.2±0.1 | 3.89±1.19 | 41.5±40.9 | 60.4±68.1 | 78±12 | 46±20 | 54±5.8 | 73±18 | 30±6.6 | 4.6±1.1 | 2.5±0.45 | 11±1.9 | 18±4.0 | 12±4.4 | 10.3±2.97 | 1.95±2.2 | 12.2±2.26 |
| 12 | 0 | 6.1±0.0 | 0.11±0.00 | 45.2±22.8 | 100 | 45 | 14 | 67 | 21 | 3.0 | 1.3 | 0.14 | 1.5 | NA | BD | NA | NA | NA |
| 12 | 15 | 6.5±0.1 | 4.70±0.40 | 19.0±1.99 | 26.9±12.1 | 77±23 | 35±3.9 | 65±7.6 | 52±5.8 | 40±22 | 5.3±0.53 | 1.4±0.26 | 20±23 | 30±34 | 13±0.61 | 7.62±5.51 | 1.55±1.22 | 9.18±6.70 |
